# Neurogenic decisions require a cell cycle independent function of the CD25B phosphatase

**DOI:** 10.1101/213074

**Authors:** Frédéric Bonnet, Mélanie Roussat, Angie Molina, Manon Azais, Sophie Belvialar, Jacques Gautrais, Fabienne Pituello, Eric Agius

## Abstract

A fundamental issue in developmental biology and in organ homeostasis is understanding the molecular mechanisms governing the balance between stem cell maintenance and differentiation into a specific lineage. Accumulating data suggest that cell cycle dynamics plays a major role in the regulation of this balance. Here we show that the G2/M cell cycle regulator CDC25B phosphatase is required in mammals to finely tune neuronal production in the neural tube. We show that in chick neural progenitors, CDC25B activity is both required and sufficient to stimulate neurogenic divisions and to promote neuronal differentiation. We design a mathematical model showing that within a limited period of time, cell cycle length modifications cannot account for changes in the ratio of the mode of division. Using a CDC25B point mutation that cannot interact with CDK, we show that part of CDC25B activity on neurogenic divisions is independent of its action on the cell cycle.

## INTRODUCTION

In multicellular organisms, managing the development, homeostasis and regeneration of tissues requires the tight control of self-renewal and differentiation of stem/progenitor cells. This issue is particularly evident in the nervous system, where generating the appropriate number of distinct classes of neurons is essential to constructing functional neuronal circuits.

Steadily increasing data reveal links between the cell cycle and stem cells' choice to proliferate or differentiate (Soufi & Dalton, 2016). The G1 phase is usually associated with the initiation of differentiation. Notably, the length of the G1 phase has been shown to play a major role in controlling cell fate decisions in neurogenesis, haematopoiesis (Lange & Calegari, 2010) and mammalian embryonic stem cells (Coronado et al., 2013; Sela, Molotski, Golan, Itskovitz-Eldor, & Soen, 2012), including human embryonic stem cells (hESCs) (Pauklin & Vallier, 2013; Sela et al., 2012). During cortical neurogenesis, a lengthening of the G1 phase is associated with the transition from neural-stem-like apical progenitors (AP) to fate restricted basal progenitors (BP) (Aral et al., 2011). Reducing G1 phase length leads to an increased progenitor pool and inhibition of neuronal differentiation, while lengthening the G1 phase promotes the opposite effects (Calegari, Haubensak, Haffner, & Huttner, 2005; Pilaz et al., 2009). In the developing spinal cord, G1 phase duration increases with neurogenesis (Kicheva et al., 2014; Saade et al., 2013). Interestingly, in hESCs and in neurogenesis it has been shown that the stem/progenitor cell uses Cyclin D, which controls G1 phase progression, to directly regulate the signaling pathways and the transcriptional program controlling cell fate choice (Bienvenu et al., 2010; Lukaszewicz & Anderson, 2011; Pauklin, Madrigal, Bertero, & Vallier, 2016; Pauklin & Vallier, 2013). A transient increase of epigenetic modifiers at developmental genes during G1 has also been reported to create “a window of opportunity” for cell fate decision in hESCs (Singh et al., 2015).

Modification of other cell cycle phases has been correlated with the choice to proliferate or differentiate. Work on hESCs reveals that cell cycle genes involved in DNA replication and G2 phase progression maintain embryonic stem cell identity (Gonzales et al., 2015), leading the authors to propose that S and G2/M mechanisms control the inhibition of pluripotency upon differentiation. In the amphibian orfish retina, the conversion of slowly dividing stem cells into fast-cycling transient amplifying progenitors with shorter G1 and G2 phases, propels them to exit the cell cycle and differentiate (Agathocleous, Locker, Harris, & Perron, 2007; Locker et al., 2006). A shortening of the S phase correlates with the transition from proliferative to differentiating (neurogenic) divisions in mouse cortical progenitors (Arai et al., 2011). In the developing spinal cord, shorter S and G2 phases are associated with the neurogenic phase (Cayuso & Marti, 2005; Kicheva et al., 2014; Le Dreau, Saade, Gutierrez-Vallejo, & Marti, 2014; Molina & Pituello, 2016; Peco et al., 2012; Saade, Gonzalez-Gobartt, Escalona, Usieto, & Marti, 2017; Saade et al., 2013; Wilcock, Swedlow, & Storey, 2007). Until now these links between cell cycle kinetics and cell fate were most often correlations, with the direct impact of cell cycle modifications on cell fate choice being only indirectly addressed. The strong correlations between the cell cycle machinery and the stem cell's choice in different model systems, emphasize the importance of elucidating how these systems work.

A link has previously been established between a regulator of the G2/M transition, the CDC25B phosphatase and neurogenesis (Gruber et al., 2011; Peco et al., 2012; Ueno, Nakajo, Watanabe, Isoda, & Sagata, 2008). The cell division cycle 25 family (CDC25) is a family of dual specificity phosphatases that catalyze the dephosphorylation of the cyclin-dependent kinases (CDKs), leading to their activation and thereby cell cycle progression (Aressy & Ducommun, 2008). Three CDC25s A, B, C have been characterized in mammals, and two, CDC25s A and B have been found in chick (Agius, Bel-Vialar, Bonnet, & Pituello, 2015; Boutros, Lobjois, & Ducommun, 2007). As observed for numerous cell cycle regulators, these molecules are tightly regulated at the transcriptional and post-transcriptional levels (Boutros et al., 2007). The N-terminal region of CDC25B contains the regulatory domain, and the C-terminal region hosts the catalytic domain and the domain of interaction with known substrates, the CDKs (Sohn et al., 2004). In Xenopus, CDC25B loss-of-function reduces the expression of neuronal differentiation markers (Ueno et al., 2008). An upregulation of CDC25B activity associated with precocious neurogenesis has been observed in an animal model of microcephaly (Gruber et al., 2011). Using the developing spinal cord as a paradigm, we previously reported that CDC25B expression correlates remarkably well with areas where neurogenesis occurs (Agius et al., 2015; Peco et al., 2012). We showed that reducing CDC25B expression in the chicken neural tube alters both cell cycle kinetics, by increasing G2-phase length, and neuron production (Agius et al., 2015; Peco et al., 2012). However, it is not clear whether the change in cell cycle kinetics is instrumental in cell fate change.

The aim of the present study is to further understand the mechanisms by which CDC25B promotes neurogenesis. First, we use a neural specific loss-of-function in mice to show that neurogenic activity of Cdc25B is conserved in mammals. Second, we use gain- and loss-of-function in chicken to show that CDC25B is necessary and sufficient to promote neuron production by controlling the mode of division. We directly measured CDC25B effect upon modes of division using recently developped biomarkers that allow to differentiate with single-cell resolution the three modes of division taking place in the developing spinal cord: proliferative where a progenitor gives rise to two progenitors (PP); asymmetric neurogenic where a progenitor gives rise to one progenitor and one neuron (PN), and terminal symmetric neurogenic where the progenitor gives rise to two neurons (NN) (Saade et al., 2013). These biomarkers were previously used to analyze the role of signaling pathways controlling the progenitor's mode of division (Le Dreau et al., 2014; Saade et al., 2017; Saade et al., 2013). CDC25B modulation of the mode of division appeared dependent on the context: in domains where cells perform mainly proliferative divisions, CDC25B gain of function promotes asymmetric neurogenic divisions, and in domains where cells accomplish mostly asymmetric neurogenic divisions, it promotes terminal symmetric neurogenic division. A mathematical model of these dynamics suggests that the cell cycle duration is not instrumental in the observed evolution of the mode of division.

Furthermore, to directly address the putative role of the cell cycle kinetics on the mode of division, we use a point mutated form of CDC25B, CDC25B^∆CDK^ unable to interact with CyclinB/CDK1 complex. We show that this molecule stimulates asymmetric neurogenic divisions and neuronal differentiation even though it does not affect the duration of the G2 phase.

## RESULTS

### Genetic Cdc25B invalidation induces a G2-phase lengthening and impedes neuron production in the mouse developing spinal cord

We previously showed that downregulating CDC25B levels using RNAi in the chicken neural tube results in a G2 phase lengthening and a reduction of the number of neurons. Here we used a genetic approach to question whether both functions are conserved in mammals, using a floxed allele of *Cdc25B* and a *NestinCre; Cdc25B+/-* mouse line to specifically ablate the phosphatase in the developing nervous system (Figure 1A). In the mouse embryo, *Cdc25B* is detected in the neural tube from E8.5 onward and remains strongly expressed in areas where neurogenesis occurs, as illustrated in the E11.5 neural tube (Figure 1B). Loss of *Cdc25B* mRNA was observed from E10.5 onward in *NestinCre; Cdc25B*^*fl/-*^ embryos (*CdcB25*^*nesKO*^ Figure 1B). We therefore determined the consequences of the Cre-mediated deletion of the floxed *Cdc25B* allele on cell cycle parameters and neurogenesis starting at E11.5.

**Figure 1.**
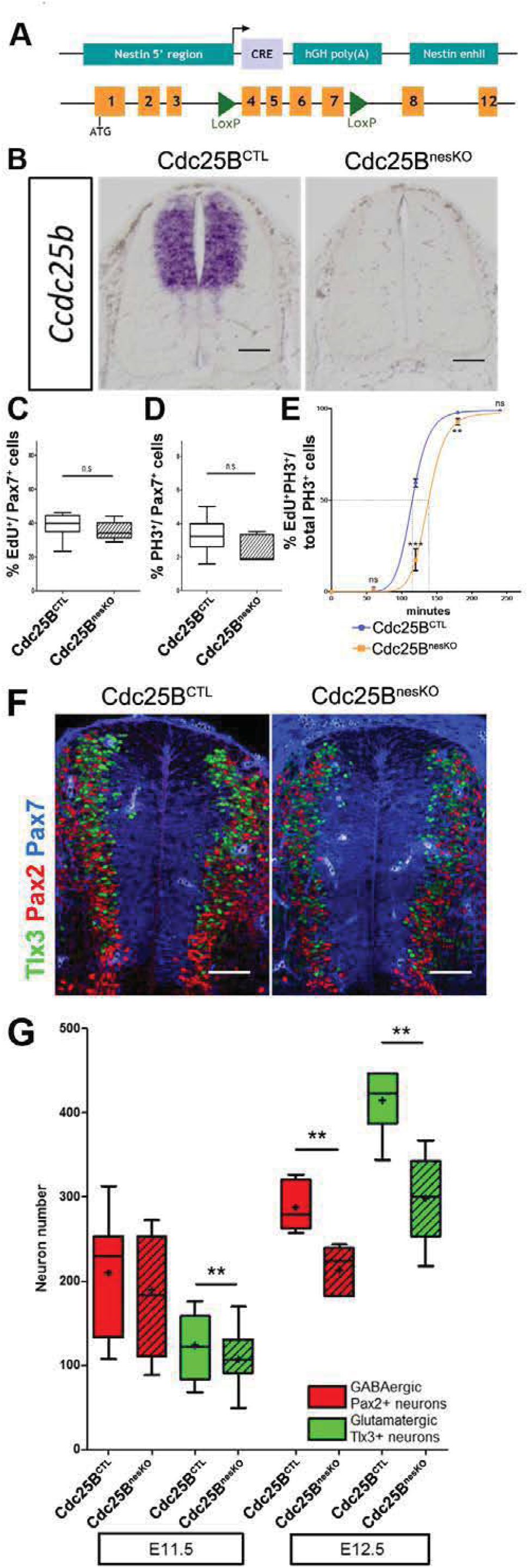
*CdclSB* conditional genetic loss-of-function increases the G2-phase length and impairs dorsal spinal neurogenesis. **A:** Scheme of the genetic construction for *Cdc25B* conditional loss-of-function. B: *Cdc25B in situ* hybridization at E11.5 in control (CTL) and nesKO conditions. C: Box and whisker plots (5/95 percentile) comparing the proliferative index: distribution of the percentage of EdU+ / Pax7+ cells indicative of the proliferative index at E11.5 in control and conditional KO neural tubes. D: Box and whisker plots (5/95 percentile) comparing the distribution of the percentage of PH3 +/ Pax7+- cells indicative of the mitotic index at E11.5 in control and conditional KO neural tubes. The proliferative index was analyzed using 20 controls and 7 nesKO embryos. E: Progression of the number of EdU/PhD corbeled nuclei with increasing EdU exposure time in control and nesKO conditions. The dashed lines correspond to 50% EdU+/PH3+ cells and indicate the G2 length. F: Cross-sections of E12.5 embryo neural tubes, stained with Pax7, Pax2 and Tlx3 in CTL and nesKO conditions. G: Box and whisker plots (5/95 percentile) comparing the distribution of the number of Pax2 and Tlx3 neurons in control and nesKO conditions at E11.5 and E12.5. The number of analyzed embryos was 15 control *vs* 11 nesKO for Pax2 and 15 control vs 10 nesKO for Tlx3. The cross indicates the mean value. Mixed model, **p<0.01. Scale bar represent 100 μm.

The proliferation capacity of the neural progenitors in *NestinCre; Cdc25B*^*fl/-*^ embryos, was determined by quantification of EdU labelled replicating neural progenitors. The proliferative index in the dorsal spinal cord (number of EdU_+_ cells among total number of neural progenitors labelled with Pax7 antibody) was similar between *NestinCre; Cdc25B*^*fl/-*^ and control embryos (*NestinCre; Cdc25B*^*fl/+*^ or *Cdc25B*^*fl/+*^ or *Cdc25B*^*fl/-*^) (Figure 1C). Similarly, the fraction of mitotic cells assessed by quantifying the number of Phospho-Histone 3 (PH3) mitotic cells in the Pax7+ cells displayed a slight and non-significant reduction in the mitotic index of mutant embryos (Figure 1D). Since downregulating *CDC25B* in the chicken neural tube resulted in a lengthening of the G2 phase, we next compared the length of the G2 phase in the dorsal spinal cord of *NestinCre;Cdc25B*^*fl/-*^ versus control embryos using the percentage of labeled mitosis (PLM) (Quastler & Sherman, 1959). Embryos were injected with EdU and allowed to recover for 1 hour, 2 hours or 3 hours before fixation and staining with EdU and PH3 antibodies. We found that the percentage of PH3/EdU positive cells is consistently lower in the dorsal domain of *NestinCre; Cdc25B*^*fl/-*^ versus control embryos (Figure 1E). The average G2-lengths extracted from the curve are 2 hours 19 minutes in mutants compared to 1 hour 49 minutes in controls (Figure 1E). This indicates that *Cdc25B* loss-of-function in dorsal neural progenitors results in a G2 phase lengthening.

The question is then whether *Cdc25B* loss-of-function affects spinal neurogenesis. Neuron production occurs in two phases in the dorsal spinal cord, an early neurogenic phase (between E9.5 and E11.5) and a late neurogenic phase (between E11.5 and E13.5) (Hernandez-Miranda, Muller, & Birchmeier, 2016). Neurons emerging from the dorsal spinal cord express numerous transcription factors including Pax2 and Tlx3 that label distinct neuron types and when combined, identify different subtypes of early (Pax2: dl4, dl6; Tlx3: dl3, dl5) and late born neurons (Pax2: dILA; Tlx3: dILB). The use of a *NestinCre* mouse line allows us to acutely ablate the phosphatase at the time of late neuron production (Hernandez-Miranda et al., 2016). We hence analyze the impact of the deletion at E11.5 and E12.5. At E11.5, the number of Tlx3+ cells is reduced in the *NestinCre;Cdc25B^fl/-^* compared to control embryos. Pax2+ neurons are also reduced yet non-significantly (Figure 1F, G). One day later, a clear and significant reduction of 25.7% and 28% in the number of Pax2+ and Tlx3+ neurons, respectively, is observed following *Cdc25B* deletion. The size of the progenitor domain measured using Pax7 immunochemistry shows a slight but non-significant increase (Figure supplement 1), indicating that neuron reduction is not due to a reduction of the progenitor population. Quantification of active caspase 3 immunostaining (E12.5) does not reveal an increase in cell death, showing that the reduction in neuron number is not due to apoptosis (not shown). The ratio of dILA to dILB neurons is similar between control (0.68) and mutant embryos (0.71), confirming that *Cdc25B* does not impact specific neuronal cell type but rather has a generic effect on neuron production. Together, these observations demonstrate that efficient spinal neuron production requires Cdc25B in mammalian embryos, illustrating that this function is conserved among higher vertebrates.

**Figure Supplement 1.**
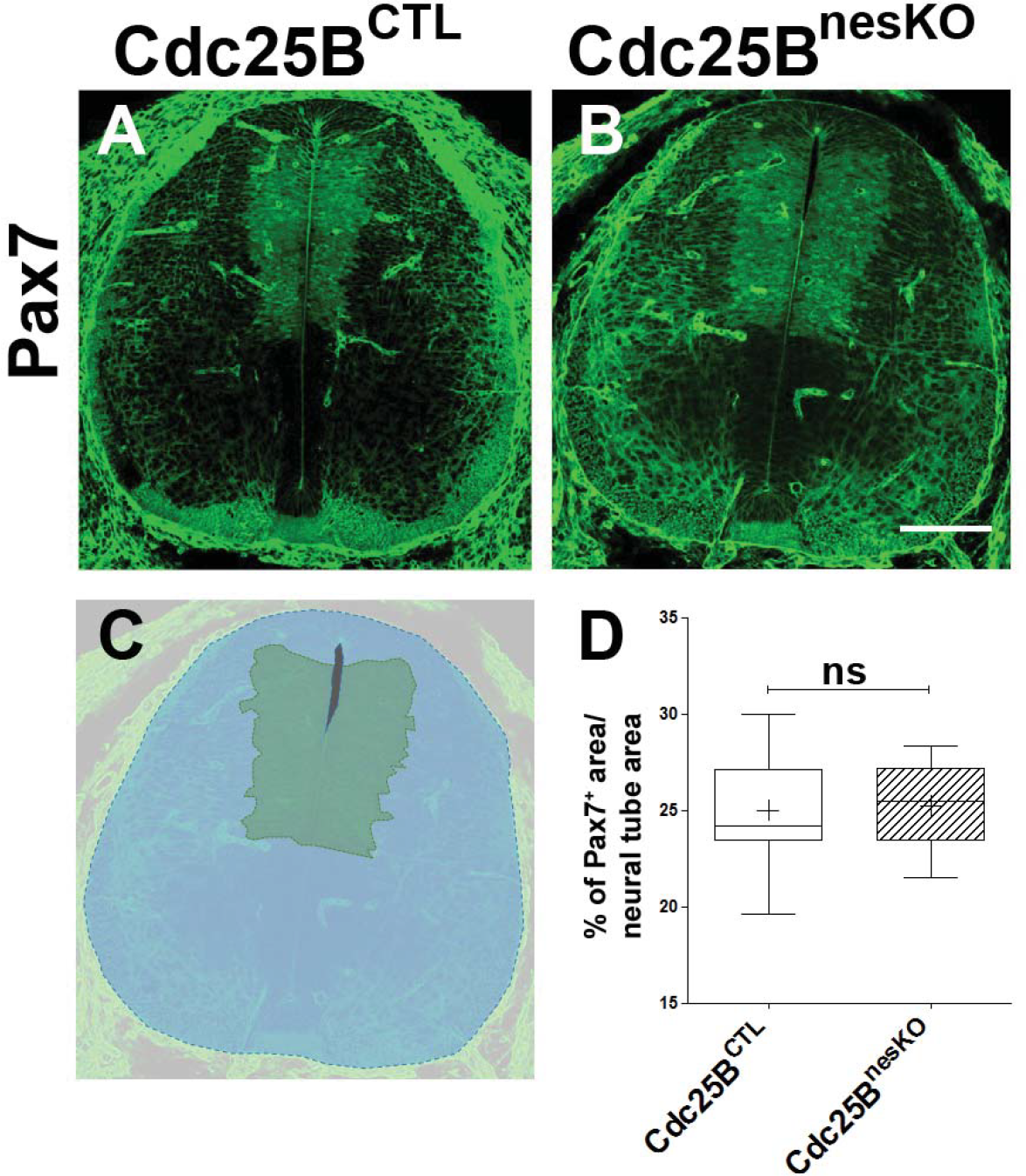
Cdc25B conditional genetic loss-of-function does not reduce the progenitor pool. **A-B**: Cross-sections of E12.5 embryo neural tubes in control (A) and conditional KO conditions (B). **C**: The progenitor pool size is evaluated by the percentage of the Pax7 progenitor area (green) compared to the neural tube area (blue). **D**: Box and whiskers plots comparing the progenitor area in a global analysis of E11.5 - E12.5 control (19 embryos) and nesKO (13 embryos) neural tubes. The cross indicates the mean value. Scale bar represents 100 μm

### CDC25B gain-of-function increases neuronal production

The fact that CDC25B downregulation impedes neuron production in mouse and chicken embryos, prompted us to test whether CDC25B gain-of-function is sufficient to stimulate neurogenesis. It is not possible to perform CDC25B gain-of-function using a robust ubiquitous promotor, because an unscheduled increase of the phosphatase during the cell cycle leads to mitotic catastrophe and subsequent apoptosis (Peco et al., 2012). To circumvent this technical impasse, we express CDC25B using the mouse cell cycle dependent CDC25B cis regulatory element (ccRE) that reproduces the cell cycle regulated transcription of CDC25B (Korner, Jerome, Schmidt, & Muller, 2001) and prevents apoptosis (Kieffer, Lorenzo, Dozier, Schmitt, & Ducommun, 2007). We verify that ccRE is sufficient to drive lacZ reporter expression in the entire chicken neural tube after transfection by *in ovo* electroporation (Figure Supplement 2A). Under the control of ccRE, the eGFP-CDC25B fusion protein is expressed in a subset of transfected cells (Figure 2A). The level of chimeric protein detected results from the periodic expression induced by the promoter and the intrinsic instability of CDC25B actively degraded at the end of mitosis. The fusion protein can be observed both in the nucleus and cytoplasm of neuroepithelial progenitors located close to the lumen (L) and in mitotic progenitors (Figure 2A arrowhead). The gain-of-function does not induce apoptosis, as revealed by quantification of active caspase 3 immunostaining (Figure Supplement 2B-D). To ascertain that the phosphatase is functional, we analyze its impact on G2 phase duration. As expected, ectopic expression of the phosphatase shortens the G2 phase (Figure 2B, blue curve) without significantly modifying the mitotic index or the proliferation index (Figure Supplement 2E-F). We analyze the neurogenic effects of CDC25B gain-of-function 48 hours after electroporation by measuring the expression of the luciferase reporter under the NeuroD promoter (Figure Supplement 3), by analyzing an interneuron marker Pax2 (Figure 2C, D) and by using a pan neuronal marker HuC/D (Figure 2F, G) in conjunction with a pan progenitor marker Sox2 (Figure 2E).

**Figure Supplement 2.**
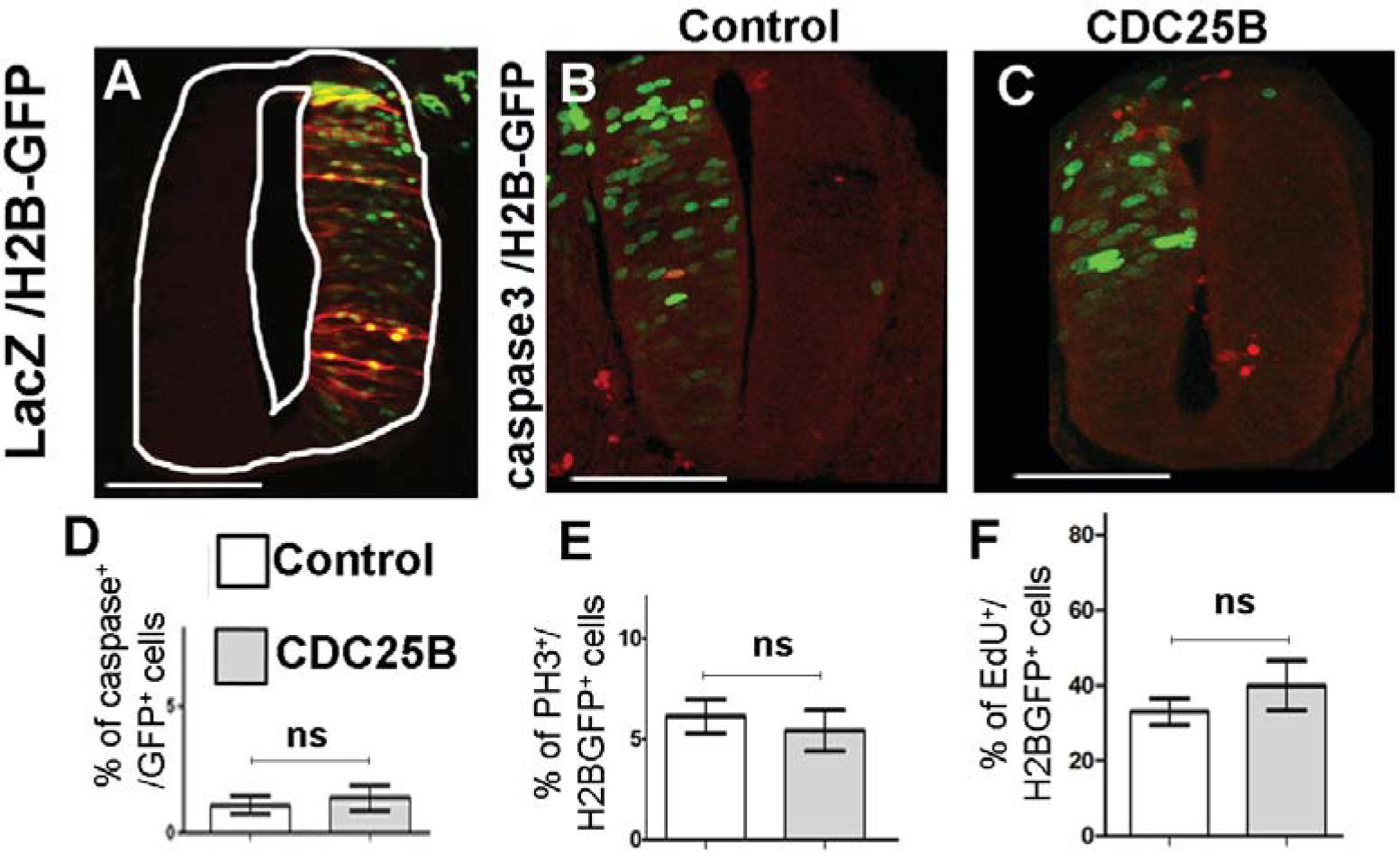
CDC25B gain-of-function does not increase apoptosis, S or M cell cycle lengths. **A**: Section of embryonic spinal cord at HH17 after co-electroporation of and pCAG::H2B-GFP and anti lacZ immunostaining in red. Note that the ccRE promoter leads to lacZ positive cells localized throughout the dorso-ventral axis of the neural tube. **B-C**: Anti active Caspase-3 immunostaining (red) 24 hours after co-electroporation of pCAG::H2B-GFP plus pccRE::lacZ (control) (B) or pccRE::CDC25B vector (C). Scale bars represent 100 μm. **D**: Percentage of active-Caspase 3^+^ cells in the H2B-GFP+ population after 24 hours: control (1.1 +/- 0.84%) and CDC25B gain-of-function (1.39 +/- 0.5%). Mean +/- sem from 3 experiments, 7 control embryos corresponding to 1194 cells, and 9 embryos corresponding to 569 cells for CDC25B gain-of-function. **E**: Mitotic index, represented as the percentage of PH3^+^ cells among H2B-GFP+ electroporated cells after 24 hours: control (6.1 +/- 0.34%) and CDC25B gain-of-function (5.4 +/- 1%). Mean +/- SEM from 3 different experiments, 8 embryos and 930 cells for the control, and 10 embryos and 868 cells for CDC25B gain-of-function. **F**: Proliferative index represented as the percentage of EdU^+^ cells in the H2B-GFP+ population after 24 hours: control (33 +/- 3.5%) and CDC25B gain-of-function (40 +/- 6.6%). Mean +/- SEM from 3 experiments, 9 embryos for the control, and 7 embryos for CDC25B gain-of-function.

**Figure 2.**
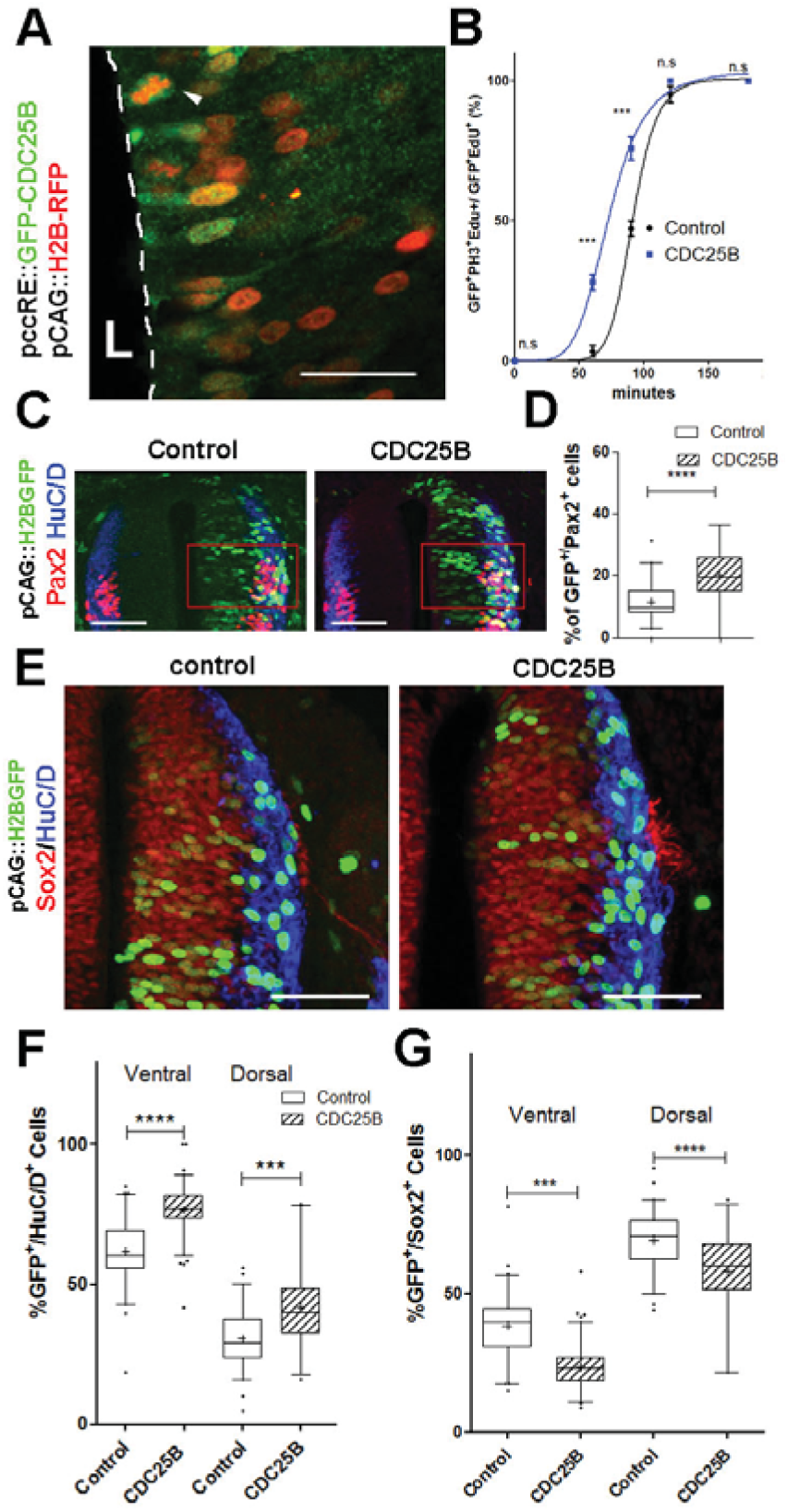
CDC25B speeds up neuronal production. **A**: Cross section of E2.5 chick spinal cord 24 hours after electroporation of pCAG::H2B-RFP vector and pccRE::GFP-CDC25B vector, followed by an anti-GFP immunolocalisation. Note that the protein is expressed in the dorsal neuroepithelium in cells exhibiting a nucleus close to the lumen side (L) or undergoing mitosis (arrowhead). Scale bar indicates 50 μm. **B**: Curves representing the progression of EdU/PH3 co-labeled nuclei with increasing EdU exposure times: control (black), CDC25B (blue). Note that the curve corresponding to the CDC25B condition (blue) is shifted to the left, showing a reduction in G2 phase length. **C**: Representative sections of E3.5 chick spinal cord 48 hours after co-electroporation of a pCAG::H2B-GFP with either a pccRE::control or a pccRE::CDC25B expression vector and processed for Pax2 (red) and HuC/D (blue) immunostaining. The red box illustrates the quantified domain. Scale bars indicate 100 μm. **D**: Box and whisker plots (5/95 percentile) comparing the percentage of Pax2^+^ cells within the electroporated population in the control and CDC25B gain-of-function experiments in the dorsal neural tube. Data from 3 different experiments with 8 embryos for the control conditions, and 5 embryos for the CDC25B gain-of-function. **E**: Representative sections of E3.5 chick spinal cord 48 hours after co-electroporation of pCAG::H2B-GFP with either a control or a CDC25B expression vector and processed for Sox2 immunostaining (red) and HuC/D (blue). Scale bars indicate 100μm. **F**: Box and whisker plots (5/95 percentile) comparing the percentage of electroporated HuC/D^+^ cells in the ventral and dorsal neural tube. Data represent 3 different experiments with 13 and 6 embryos in dorsal and ventral respectively under control conditions and 6 and 7 embryos in dorsal and ventral respectively for CDC25B gain-of-function. The cross represents the mean value. **G**: Box and whisker plots (5/95 percentile) comparing the percentage of Sox2^+^ cells within the electroporated population in the control, CDC25B gain-of-function experiments in the dorsal or ventral neural tube. Same conditions as in F.

A quantitative analysis performed on the entire neural tube using NeuroD- reporter assay indicates that increasing CDC25B is sufficient to promote neuronal commitment (Figure Supplement 3). In the neural tube, development of the ventral progenitor population is usually considered more advanced than its dorsal counterpart (Kicheva et al., 2014; Saade et al., 2013). Accordingly, the temporality of neuron production progresses from ventral to dorsal (Kicheva et al., 2014; Saade et al., 2013) and correlated with endogenous CDC25B expression (Peco et al., 2012). We therefore analyze separately the fraction of neurons generated following CDC25B gain-of-function in the ventral and dorsal halves of this structure. In the ventral neural tube, CDC25B gain-of-function increases the percentage of HuC/D^+^ GFP^+^ cells from 61.6 +/- 1.5% to 76.5 +/-0.9 %. Similarly, in the dorsal spinal cord, the proportion increases from 30.66+/- 1.34% to 41.80+/-2.64% with the CDC25B gain-of-function (Figure 2F, G). A significant increase in neurogenesis is also observed using Pax2 immunostaining from 11.4 +/- 1 % to 20 +/-1.8 % (Figure 2C, D). Conversely, CDC25B gain-of-function reduces the proportion of cells expressing the progenitor marker Sox2 (Figure 2E). Together, these results indicate that CDC25B is sufficient to stimulate neuron production.

**Figure Supplement 3.**
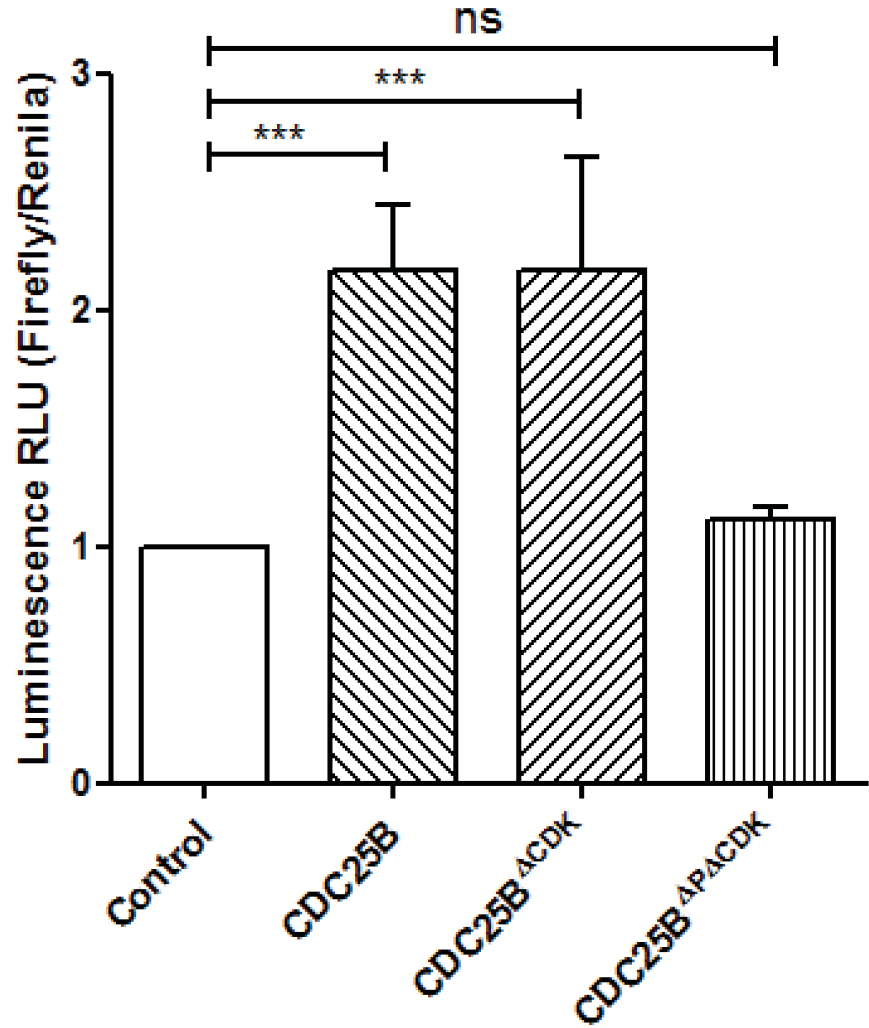
Effects of various CDC25B constructs on NeuroD promoter activity. Column bar graph representing the transcriptional activity of the NeuroD promoter assessed *in vivo* following electroporation of the indicated CDC25B constructs. At HH11 the embryos were electroporated with the pNeuroD::Luc vector and a renilla luciferase reporter construct carrying the cytomegalovirus immediate early enhancer promoter for normalization (Promega), together with the indicated DNAs. At HH22, 48 hours post electroporation, the neural tubes were dissected and processed following the Dual Luciferase Reporter Assay System protocol (Promega). The data are presented as the means ± SEM from at least 14 embryos in 4 experiments.

### CDC25B has no effects on mitotic spindle parameters

An increase in CDC25B activity has been shown to induce a shifted cleavage plane and precocious neurogenesis during corticogenesis in mouse (Gruber et al., 2011). We therefore tested the effect of CDC25B gain-of-function on spindle orientation in spinal neural precursors. We measured the angle of mitotic spindle as previously described (Saadaoui et al., 2014). We did not observe a significant change in the spindle orientation (Figure 3A, B). Another element implicated in asymmetric cell fate in neural progenitors is the spindle size asymmetry (SSA), i.e., the difference in size between the two sides of the spindle (Delaunay, Cortay, Patti, Knoblauch, & Dehay, 2014). Our CDC25B gain-of-function experiments did not induce a significant modification of the SSA of chick spinal neural progenitors (Figure 3C-D). In summary, our analyses did not reveal an effect of CDC25B activity on the orientation or the size of the mitotic spindle.

**Figure 3.**
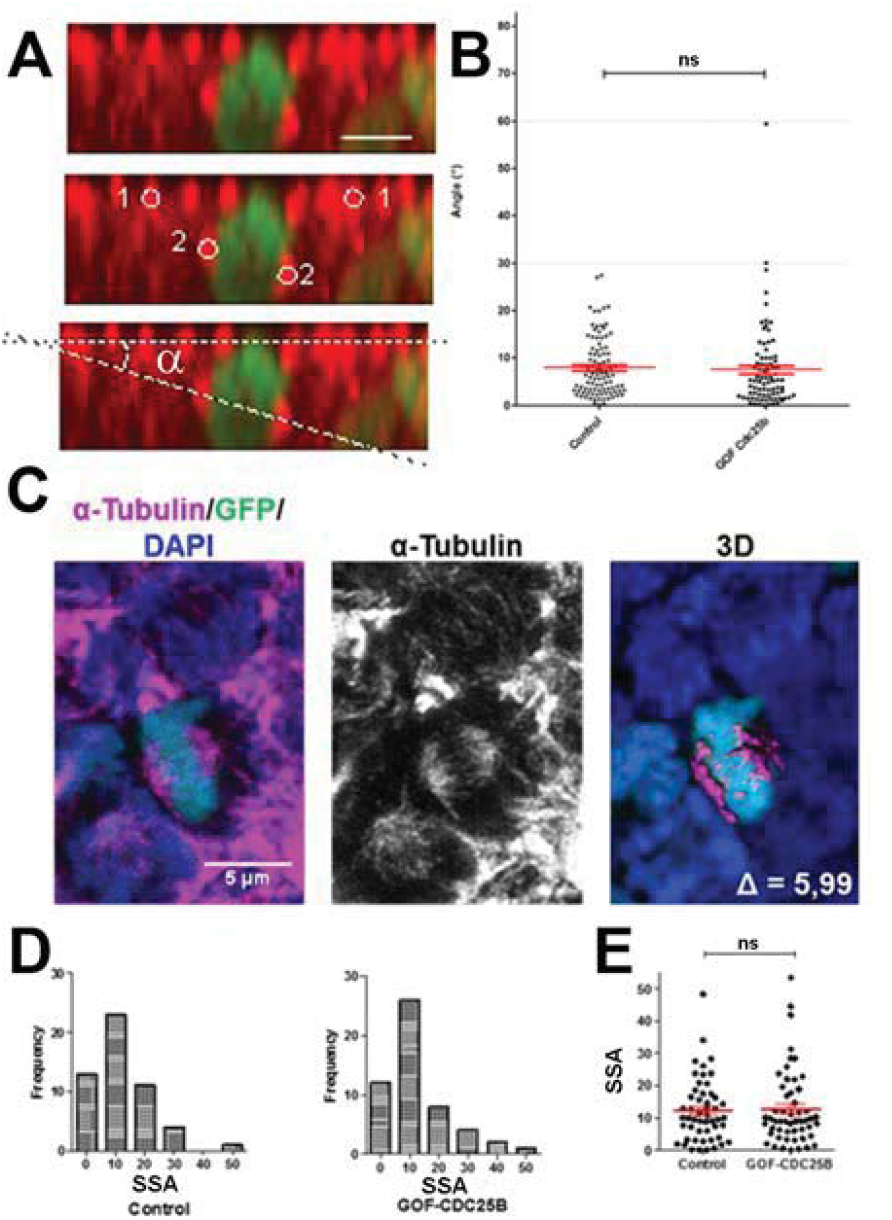
CDC25B ga in-of-f unction does not affect mitotic spindle orientation or spindle-size asymmetry (SSA). **A**: Representative Z plane image of an anaphase cell expressing H2B-GFP that decorates chromosomes (green) and immunostained with *γ* tubulin to label centnosomes (red). Aligned interphase centnosomes labelled as 1 and mitotic spindle poles labelled as 2 (middle image) were used to measure mitotic spindle angle *α* (lower image). Scales bar represent 5 Mm. **B**: Quantification of mitotic spindle angle *α*, 24 hours after electroporation in control and CDC25B gain-of-function experiments. C: Representative image of a symmetric metaphase cell: H2B-GFP and DAPI stain the nuclei and *α*-tubulin stains the mitotic spindle (left and middle images). Right image, 3D reconstruction of the symmetric spindle using Imaris which measures the spindle-size delta. D. E: Distribution of the Spindle Size Asymmetry (SSA) size difference between the two sides of the spindle 24 hours after electroporation: Box plot of the SSA distribution (D) and scatter plot of SSA distribution (E).

### CDC25B downregulation maintains proliferative divisions at the expense of both asymmetric and symmetric neurogenic divisions

To elucidate CDC25B function, we investigate whether it might promote neurogenesis by controlling the division mode of neural progenitors. We take advantage of a strategy recently developed by E. Marti and colleagues (Le Dreau et al., 2014; Saade et al., 2017; Saade et al., 2013), which allows us to unequivocally identify and distinguish the three modes of division, PP, PN and NN, occurring in the chicken developing spinal cord. Briefly, neural tube is electroporated with the Sox2::GFP and Tis21::RFP reporters, and 24 hours later the number of neural progenitors expressing each of these markers is quantified at mitosis. Thus, cells performing PP divisions express only Sox2::GFP and appear in green, those performing NN divisions express only Tis21::RFP and appear in red, while asymmetric neurogenic divisions, PN, which co-express both biosensors, appear in yellow (Figure 4A). Using these biomarkers in the dorsal neural tube, we obtained a number of PP, PN and NN divisions comparable to the ones previously described (Figure 2B) (Le Dreau et al., 2014). Because the number of electroporated cells in mitosis is very small, we determine whether counting neural progenitors displaying green, yellow or red fluorescence is equivalent to counting only mitotic cells in the dorsal spinal cord 24 hours post electroporation. We do not detect a significant difference in the % of green (GFP+), yellow (GFP+/RFP+) and red (RFP+) cells in total neuroepithelial progenitors (55.4 +/-6.2% green cells, 29.3 +/- 3.9% yellow cells and 15.2 +/- 2.9% red cells) and during mitosis (57.9 +/- 9.3% green cells, 23.2 +/- 8.5% yellow cells and 19 +/- 7.3% red cells) (Figure 4B). We therefore use the percentage of labeled progeny to estimate the percentage of proliferative (PP), asymmetric neurogenic (PN) and terminal neurogenic (NN) divisions. Because of reporter stability, the temporal window of analysis of Marti's biosensors is restricted to 24 hours (Saade et al., 2013). CDC25B RNAi electroporation leads to a consistent and strong downregulation in *CDC25B* transcripts located in the intermediate neural tube (Figure 4C bracket). We therefore determine the impact of *CDC25B* downregulation on the mode of division in progenitors located in this domain. We co-electroporate the biomarkers with either the CDC25B-RNAI plasmid, or the control scrambled plasmid at stage HH 11 and quantified the number of green (PP), yellow (PN) and red (NN) cells 24 hours later at stage HH 17 (Figure 4D-E). When compared to the control scrambled RNAi, the CDC25B RNAi induces a massive increase in green PP progeny (13.4 ± 1.31 % to 35.1 ± 1.82%), mostly at the expense of yellow PN progeny (from 72.1 ± 1.85% to 56.2 ± 1.70 % and to some extent, of the red NN progeny (from 14.6 ± 1.43% to 8.74 ±0.8%, Figure 4E).

**Figure 4.**
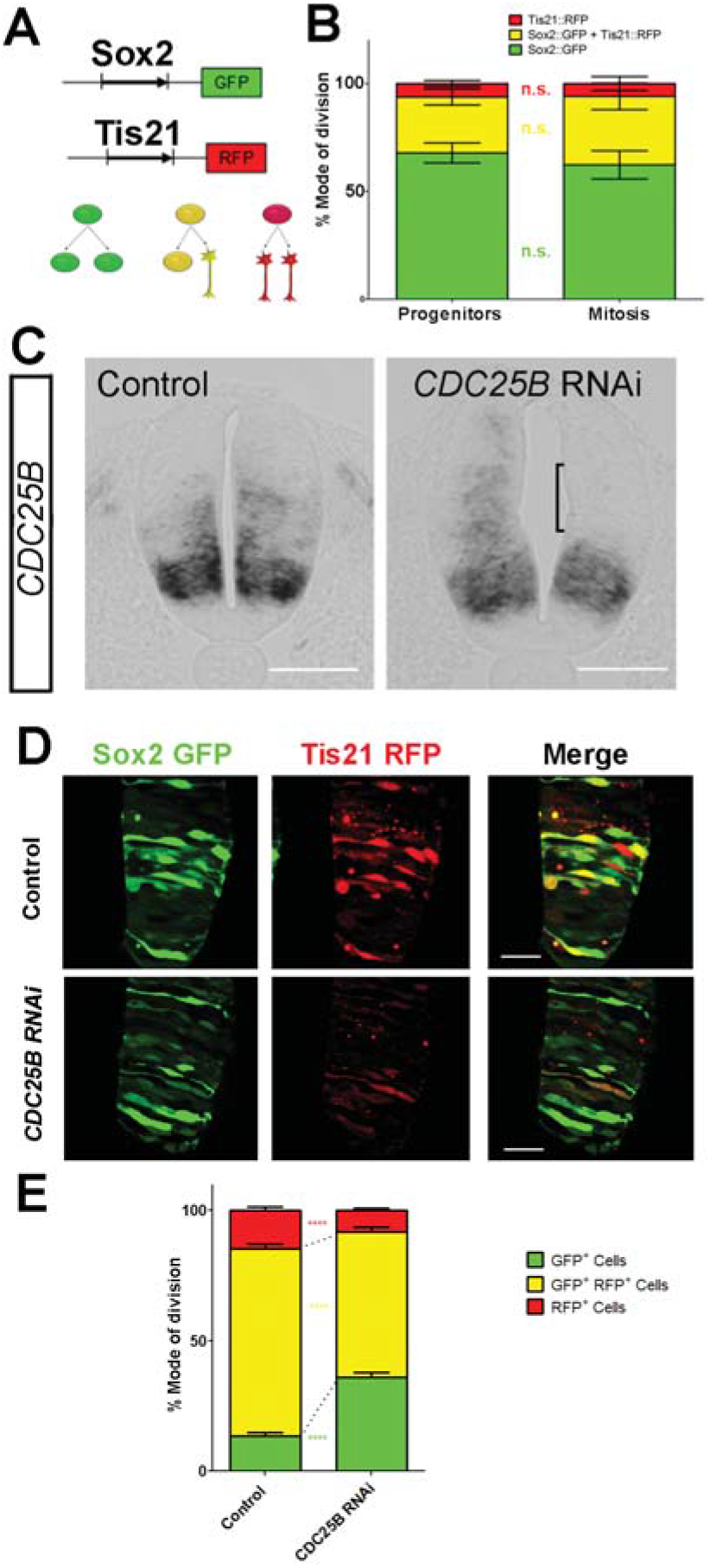
CDC25B downregulation reduces neurogenic divisions. **A**: Schematic representation of the Sox2::GFP Tis21::RFP labelling strategy. A GFP expressing cell (green cell) corresponds to a PP division, a cell expressing both GFP and RFP (yellow cell) corresponds to a PN division, and a RFP expressing cell (red cell) corresponds to a NN division. **B**: Histograms representing the percentage of cells expressing the reporters Sox2::GFP and Tis21::RFP at HH17 in the entire progenitor’s population or in progenitors performing mitosis identified with phospho-histone-3 (PH3) immunostaining. Note that these results are not significantly different. These data are obtained from 3 different experiments, 7 embryos, 365 progenitors, and 79 mitoses. **C**: In situ hybridization for CDC25B on HH17 spinal cord, 24 hours post electroporation of Control RNAi (left panel) and CDC25B RNAi (right panel). The reduction of CDC25B expression in the intermediate region is indicated by a bracket. Cells were electroporated on the right side of the neural tube (not shown). Scale bars indicate 100 μm. **D**: Cross-sections of chick spinal cord at HH17, 24 hours after co-electroporation of Sox2p::GFP and Tis21p::RFP reporter, plus a control RNAi vector or the CDC25B-RNAi vector. Scale bars indicate 50 μm. **E**: Histograms representing the percentage of progenitors expressing Sox2p::GFP and Tis21p::RFP 24hrs after co-electroporation of a control vector or a CDC25B RNAi vector. 4 experiments include 7 control embryos and 15 *CDC25B* RNAi embryos.

This observation indicates that CDC25B downregulation hindered neuron production by maintaining proliferative divisions at the expense of asymmetric and symmetric neurogenic divisions.

### CDC25B Gain-of-function promotes asymmetric and symmetric neurogenic divisions

We then use the same strategy to test how CDC25B gain-of-function affects the mode of division. At the time of electroporation (stage HH11), the neural tube contains essentially self-expanding progenitors (Le Dreau et al., 2014; Saade et al., 2013). 24 hours later, (stage HH17), the repartition of the modes of division is not the same in dorsal and ventral control conditions. Dorsal neural tube contains mainly self-expanding progenitors (66.3% Sox2^+^cells, Figure 5A, B) and (Le Dreau et al., 2014), whereas ventral neural tube encloses essentially neurogenic progeny (61.7% of Sox2VTis21^+^ cells, Figure 5A, B) and (Saade et al., 2013), in accordance with the temporality of neurogenesis which progresses from ventral to dorsal.

**Figure 5.**
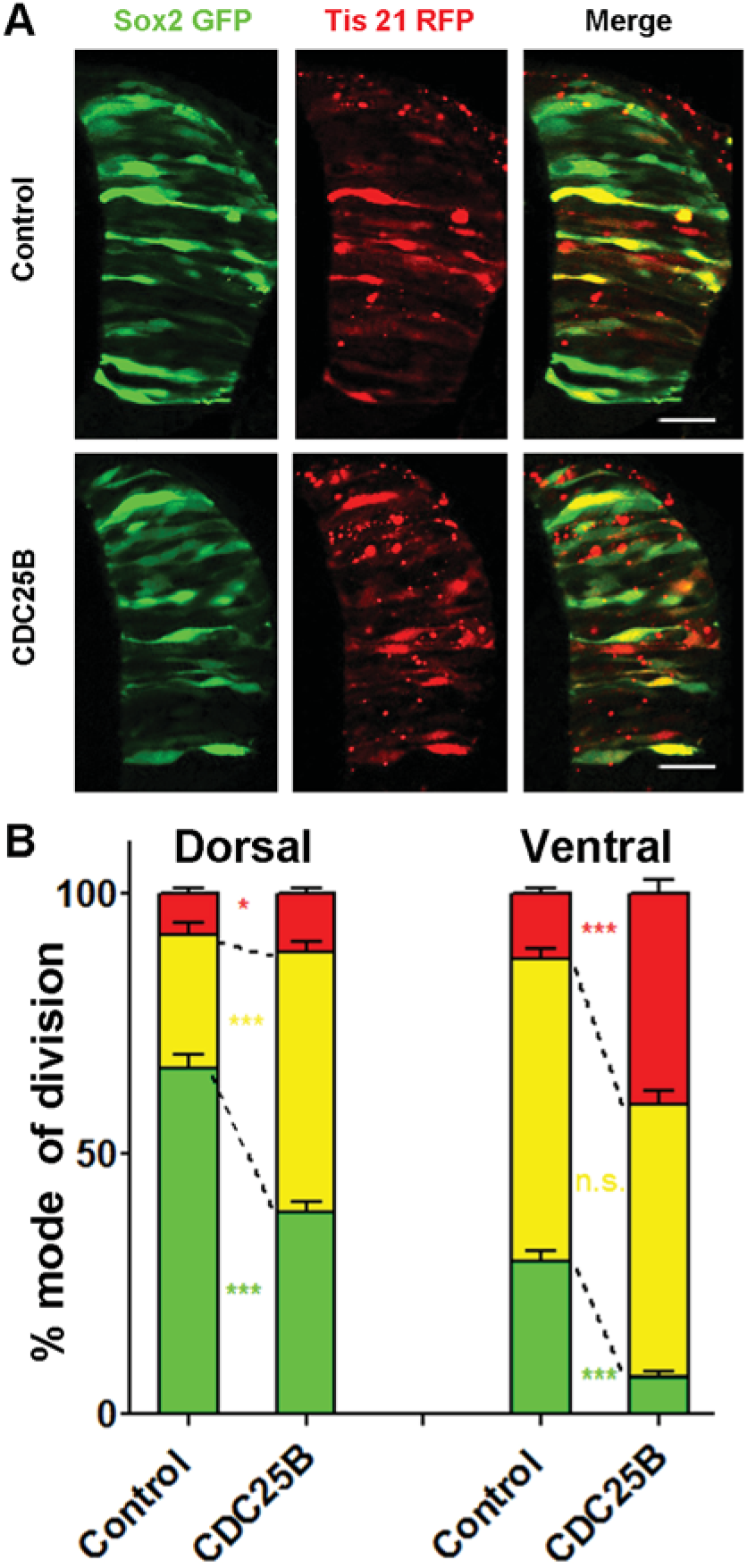
CDC25B gain-of-function promotes neurogenic divisions. **A**: Representative cross-sections of HH17 chick spinal cord, 24 hours after electroporating Sox2p::GFP and Tis21p::RFP reporters, plus a control vector pccRE::lacZ, or a pccRE::CDC25B vector. Scale bars indicate 50 μm. **B**: Histograms representing the percentage of progenitors expressing Sox2p::GFP and Tis21p::RFP 24 hours after co-electroporation with control or CDC25B vectors in the dorsal and ventral spinal cord. Data represent the means +/- sem. Data represent 3 different experiments with 5 and 10 embryos in dorsal and ventral respectively under control condition and 5 and 6 embryos in dorsal and ventral respectively under CDC25B gain-of-function condition.

In the dorsal neural tube, CDC25B gain-of-function leads to a reduction in the percentage of PP progeny (from 66.3 ± 2.6 to 38.6 ± 2.1%) and a concomitant, increase in the percentage of PN neurogenic progeny (from 25.9 ± 2.1 to 50.1 ± 1.9%). In this tissue, the percentage of NN progeny progresses only slightly (from 7.8 ± 1.2 to 11.3 ± 1 %, Figure 5B). This observation indicates that CDC25B gain-of-function in early steps of neurogenesis reduces proliferative divisions and increases asymmetric neurogenic divisions.

In the ventral neural tube, CDC25B gain-of-function induces a massive reduction of proliferative progeny (from 39.3 +/- 1.3% to 6.9 +/- 1%) and led to an increase in NN progeny (from 12.7 +/- 1.1% to 40.7 +/- 2.7%), without significantly modifying the percentage of PN cells (from 58 +/- 2% to 52.3 +/- 2.8%, Figure 5B). Thus, CDC25B ectopic expression in a more advanced neural tissue reduces proliferative divisions and increases terminal neurogenic divisions.

Together, these results suggest that CDC25B activity in neural progenitors reduces proliferative divisions, promoting either asymmetric or symmetric neurogenic divisions, depending on the receiving neural tissue.

### Mathematical modelling reveals that cell cycle duration is not instrumental in controling the mode of division

To test quantitatively our data from a dynamical point of view, we formalize in mathematical terms, the current understanding of what happens in this biological system (Figure 6A). We consider a population of progenitors, P(to), at time to, and we assume that their different modes of division result in expanding either the pool of progenitors P(t) through proliferative divisions (PP divisions) or the pool of neurons N(t) by neurogenic divisions (PN and NN divisions).

**Figure 6.**
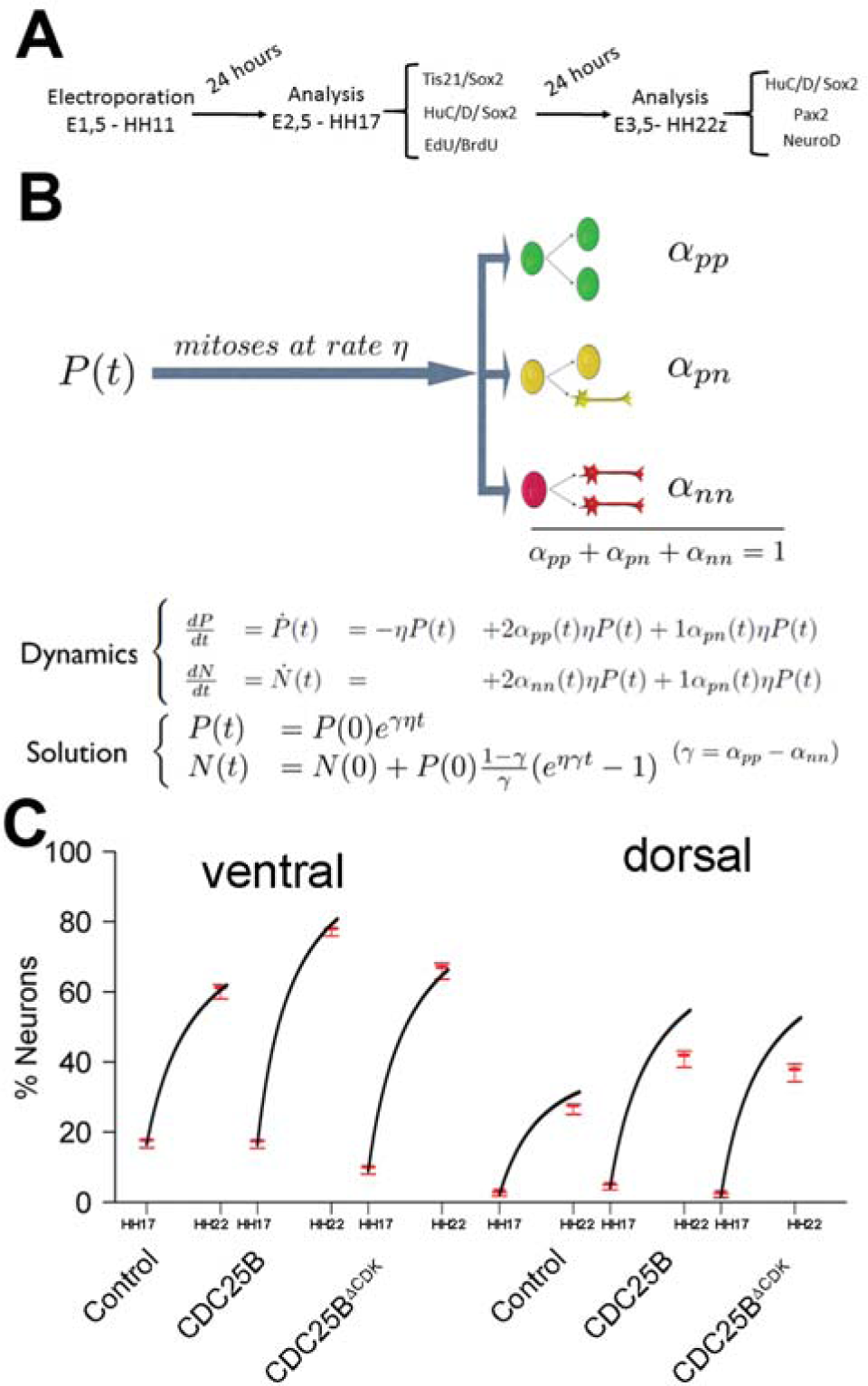
mathematical model linking the mode of division to the fraction of neurons generated. **A**: Scheme of the experimental time course. Neural tubes are electroporated at stage HH11. 24 hours (HH17) and 48 hours (HH 22) post electroporation cell cycle parameters, mode of division and progenitor/neuronal markers are analyzed. **B**: Illustration of our mathematical model. We consider P(t) a pool of progenitors at a given time with a mitotic rate η. These mitoses lead up to three kinds of mode of division: a fraction α_pp_ producing symmetric proliferative divisions yielding two progenitors, a fraction α_pn_ producing asymmetric divisions yielding one progenitor and one neuron (a precursor of), and a fraction α_nn_ producing symmetric neurogenic divisions yielding two neurons. The equations display the dynamics governing the pools of progenitors P(t) and neurons N(t) at any time t. These dynamics are solved for a given initial condition P(0), N(0), and we obtained the state of the system any time later (Solution, details in Supplement information text 1). **C**: Predictions of the kinetics of the neuronal fraction between stage HH 17 and 22 in the different conditions, compared to the mean +/- confidence interval 95% (in red) of the experimental data at stage HH17 and HH22 (from Figure 2F and 7C).

Denoting η, the rate at which P cells undergo divisions per unit time (which depends only on the cell cycle duration), the growth rates of the two pools only depend on the relative magnitude of each mode of division.

Denoting α_PP_, α_PN_ and α_NN_ the corresponding proportions of the modes of division (their sum is 1), the growth rates of the two pools (i.e. their time derivatives P(t) and N(t) for Progenitors and Neurons respectively) can then be directly formalized as:

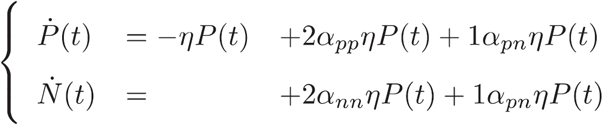

In this model, the evolution of the pool of progenitors is governed by α_PP_ and α_NN_ (because α_PN_ does not affect the pool of progenitors, only the pool of neurons). Denoting *γ* = α_pp_ - α_NN_ the difference between the two proportions, we then have that *γ*=1 (α_PP_= 1, α_NN_ =0) corresponding to purely self-expanding progenitors and *γ* =-1 (α_PP_=0, α_NN_ =1) corresponding to fully self-consuming progenitors. Hence *γ* is a good indicator of the balance between proliferation and differentiation of the progenitors.

Using *γ*, the model can be rewritten more simply as:

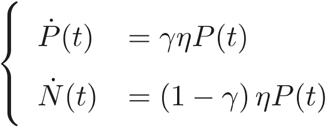

An explicit solution is:

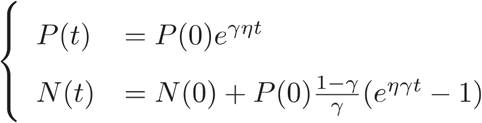

This equation means that if the quantities of progenitors and neurons are determined at a given time t (P(0), N(0)), e.g. at HH17, we can compute the expected number of progenitors and neurons at any time later, e.g. at HH22, provided that the modes of division and cell cycle times can be considered constant over the considered period. Full details of the mathematical work are given in Supplement Information Text 1.

We then compare quantitatively the experimental data to the predictions based on our current hypotheses. This comparison is surprisingly auspicious for the control and gain-of-function experiments in the ventral zone (Figure. 6C, left). In this zone, considering the ratio between the two pools at HH17 (e.g. the measured fractions of neurons), the measured cell cycle duration (12 hours), the set of modes of division measured at HH17, and the hypothesis that those modes of divisions keep unmodified during 24 hours, the model predicts with good accuracy the ratios between the two pools at HH22. In the dorsal zone, the model correctly predicts the control condition, and it confirms the tendency of CDC25B gain-of-function to promote a greater neuron fraction, albeit with some quantitative discrepancy (the model overestimates the fraction of neurons). This suggests that, notwithstanding biological complexity, the general picture of a pool of progenitors among which cells undergo different modes of division, appears relevant.

Our model is built on the assumption that all cells undergo mitosis at the same rate, and that the fate of any mitosis is stochastic and probabilistically distributed according to the fraction of dividing cells undergoing PP, PN or NN divisions, a common division rate for all progenitors associated with probabilistic fates (Supplement Information Text 1 paragraph 3.1). In this picture, the proportion of mode of division controls directly the numbers of progenitors and neurons that are generated. However, the model is compatible with an alternative interpretation, in which the three modes of division correspond to specific division rates associated with deterministic fates (Supplement Information Text 1, paragraph 3.2). In this case, each population of progenitors has a specific mean cycling time and the cell cycle time is instrumental to the mode of division. Namely, cycling at rate 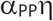 would result in a PP division, cycling at rate 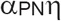 would result in a PN division, and cycling at rate 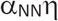 would result in a NN division. Therefore, the numbers and proportions of progenitors / neurons at HH22 would result from the difference between cell cycle times associated with modes of division. We compute these putative cell cycle times based on the data obtained in the three conditions and the two zones (Table 1). The wide range of specific cycle times, i.e., from 17 to 172.7 hours, is incompatible with data usually recorded (reviewed in (Molina & Pituello, 2016)). This suggests that, in the time window of our analyses, the observed evolution of progenitors and neurons cannot be directly explained by limited differences in cell cycle durations among the three modes of division.

**Table 1.** Putative time it would take to achieve the three kinds of division under a model which assumes that only cycle time determines the fate output. Full consequences derived from this assumption are given in Supplement Information Text 1, section 3.2. Basically, such an assumption would imply that cycling rates associated with each mode of division should be proportional to the observed fraction of that mode. If we observe, for instance, 60% PP-divisions and 10% NN-divisions (like it is about the case in the Control dorsal), then a NN-division should take 6 times as long as a PP-division. If we exclude such a possibility, then the distribution of fates cannot be exclusively determined by differences in fate-based cycle times. It does not exclude that a given kind of fate (e.g. proliferative divisions PP) could require a longer time to be achieved than others, it excludes that such differences would suffice per se to explain the differences between the fractions of fates.

|  | T<sub>PP</sub> (hours) | T<sub>PN</sub> (hours) | T<sub>NN</sub> (hours) | T<sub>c</sub> (hours) |
| --- | --- | --- | --- | --- |
| Control dorsal neural tube | 18,1 | 46,3 | 154,1 | 12,0 |
| CDC25B dorsal neural tube | 31,1 | 23,9 | 106,0 | 12,0 |
| CDC25B<sup>Δcdk</sup> dorsal neural tube | 29,8 | 23,5 | 150,0 | 12,1 |
| Control ventral neural tube | 41,0 | 20,7 | 94,5 | 12,0 |
| CDC25B ventral neural tube | 172,7 | 22,9 | 29,5 | 12,0 |
| CDC25B<sup>Δcdk</sup> ventral neural tube | 72,2 | 17,0 | 94,7 | 12,0 |

### CDC25B acts on asymmetric neurogenic division independently of CDK interaction

One prediction of our model is that neurogenesis might be affected independently of cell cycle length modification. To test whether the CDC25B-induced G2 phase modification is instrumental in promoting neurogenesis, we use a mutated form of CDC25B that was shown not to affect cell cycle kinetics. The mutation prevents CDC25B-CDK1 interactions without affecting CDC25B phosphatase activity (Sohn et al., 2004). Accordingly, expressing this mutated form of the phosphatase called CDC25B^∆CDK^ does not modify G2 phase length in neuroepithelial progenitors (Figure 7A, red curve). 24 hours after electroporation of CDC25B^∆CDK^ in the dorsal neural tube, we observe a reduction of PP progeny (from 66.3 +/-2.7% to 40.2 +/- 2.5%), an increase in PN progeny (from 25.9 +/-2.1% to 51.1 +/-2.2%), and no effect in NN progeny (from 7.8 +/-1.2% to 8.0 +/-1.1%, Figure 7B). In this context, the fraction of HuC/D^+^ neurons generated 48 hours following CDC25B^∆CDK^ expression increases from 30.7 +/- 1.3% to 40.4 +/- 2.5%. (Figure 7C). Similarly, the percentage of Pax2^+^neurons is increased from 11.3 +/-1 % to 18.3 +/-1.3% (Figure 7D).

**Figure 7.**
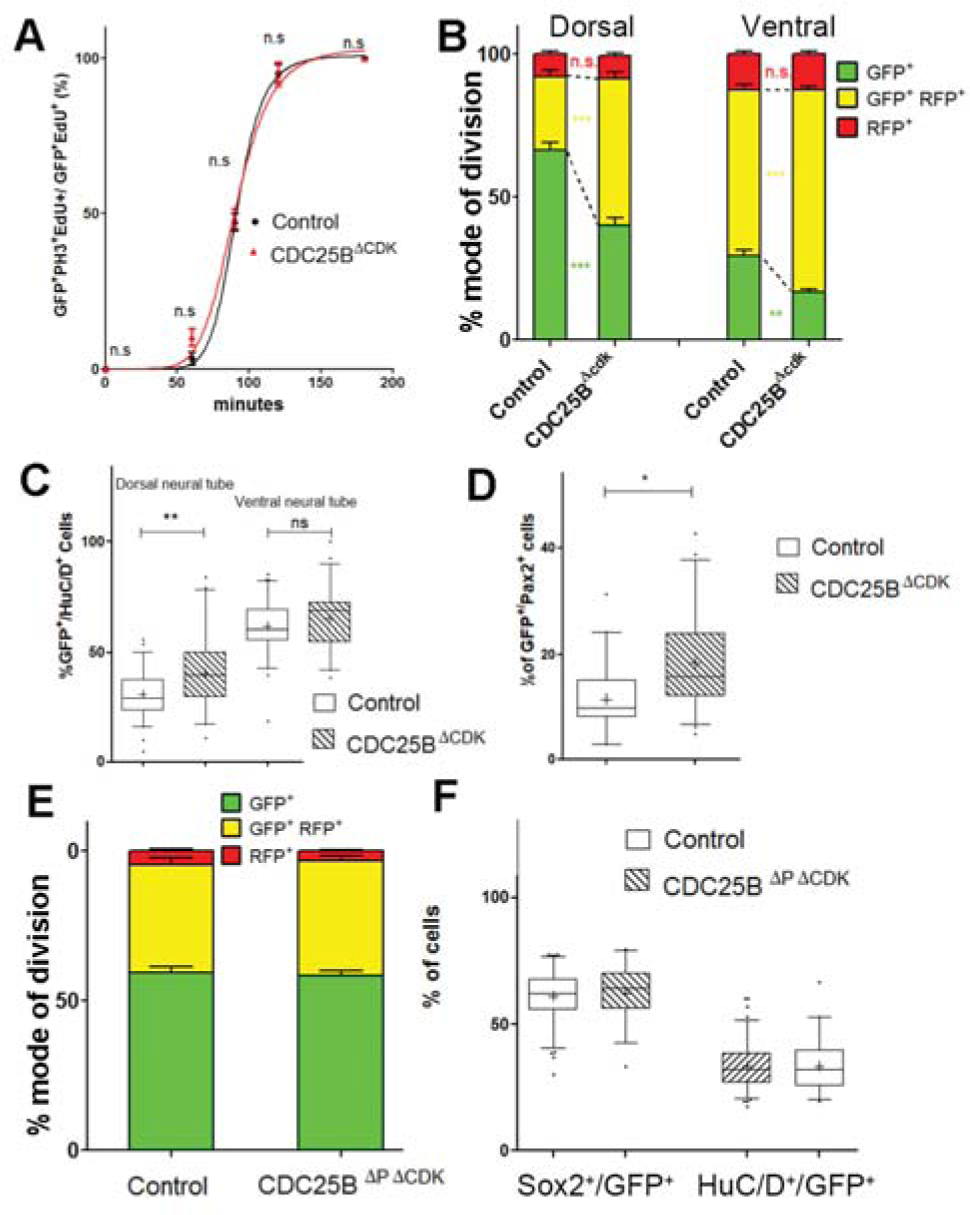
CDC25B gain-of-function promotes neurogenesis independently of CDK interaction. **A**: Curves representing the progression of EdU/PH3 co-labeled nuclei with increasing EdU exposure times: control (black), CDC25B^∆CDK^ (red). Note that the curve for the CDC25B^∆CDK^ condition is similar to the control, indicating an absence of effect on G2 length. **B**: Histograms representing the percentage cells expressing Sox2p::GFP and Tis21p::RFP 24 hours after co-electroporation with control or CDC25B^∆CDK^ vectors in the dorsal or ventral spinal cord Data represent the means +/- sem. Data represent 3 different experiments with 5 and 10 embryos in dorsal and ventral respectively under control conditions and 4 and 9 embryos in dorsal and ventral respectively for CDC25B^∆CDK^ gain-of-function. **C:** Box and whisker plots (5/95 percentile) comparing the percentage of HuC/D+ cells within the electroporated population in control or CDC25B^∆CDK^ gain-of-function experiments in the dorsal or ventral neural tube at HH22. Data represent 3 different experiments with 13 and 6 embryos in dorsal and ventral respectively under control conditions and 6 and 3 embryos in dorsal and ventral respectively for CDC25B^∆CDK^ gain-of-function. **D:** Box and whisker plots (5/95 percentile) comparing the percentage of Pax2 positive cells in the dorsal neural tube at HH22. Data from 3 different experiments with 8 embryos for control conditions, and 11 embryos for CDC25B^∆CDK^ gain-of-function. The cross represents the mean value. **E:** Histograms representing the percentage of progenitors expressing Sox2p::GFP and Tis21p::RFP at HH17, 24h after electroporation of a control or CDC25B^∆CDK∆P^ expressing vector in the dorsal half of the spinal cord. Data from 3 different experiments with 6 embryos for the control, and 9 embryos CDC25B^∆P∆CDK^. **F:** Box and whisker plots (5/95 percentile) comparing the percentage of Sox2^+^ or HuC/D^+^ cells within the electroporated population in the control or CDC25B^∆P∆CDK^ gain-of-function experiments in the dorsal spinal cord at HH17. Data from 3 different experiments with 11 embryos under control conditions and 6 embryos for CDC25B^∆P∆CDK^ The cross indicates the mean value.

In the ventral neural tube, CDC25B^∆CDK^ overexpression leads to a reduction of PP progeny (29.3 +/- 2.1% vs 16.6 +/- 1.2%), an increase in PN progeny (58 +/- 2% vs 70.7 +/-1.4%) and no effect on NN progeny (12.7 +/-1.1% vs 12.7 +/-1.1%, Figure 7B). In both ventral and dorsal domains, the CDK mutated form promotes asymmetric neurogenic divisions but is not able to promote terminal symmetric ones. In accordance with the effects on the mode of division, in the ventral neural tube, CDC25B^∆CDK^ induces a slight but non-significant increase of HuC/D expression (Figure 7C). We take advantage of our mathematical model to determine whether this slight increase in neuron production is coherent with the fact that the mutated form does not promote NN divisions, and the number of neurons predicted is in agreement with the experimental data (Figure 6C). To determine whether CDC25B^∆CDK^ function on asymmetric division and neuronal differentiation requires phosphatase activity, we use a form of the protein containing an additional point mutation inactivating the catalytic domain (CDC25B^∆P∆CDK^). This construct does not affect the mode of division at 24 hours (Figure 7E). 48 hours post electroporation this mutated form does not modify NeuroD reporter expression (Supplement Figure 3), the percentage of HuC/D^+^ neurons or the percentage of Sox2^+^ progenitors populations (Figure 7F), indicating that the phosphatase activity is required for the neurogenic function of CDC25B.

Altogether, these results show that the CDC25B phosphatase is necessary and sufficient to promote neurogenesis via a modification of the mode of division. Importantly, CDC25B^∆CDK^ stimulates asymmetric neurogenic divisions and neuronal differentiation without affecting the duration of the G2 phase. This opens the possibility that the phosphatase possesses a cell cycle independent neurogenic function.

## DISCUSSION

An important issue in the field of neurogenesis concerns the implication of cell cycle function during neuron production (Agius et al., 2015). Here, we confirm in mammals our previous observations in birds, that the G2/M cell cycle regulator CDC25B phosphatase is required to finely tune neuronal production in the neural tube. Gain-of-function experiments performed in the chick neural tube reveal that CDC25B activity is sufficient to modify the mode of division of neural progenitors and to promote neuronal differentiation concomitantly with a shortening of the G2 phase length. We demonstrate that CDC25B expression in neural progenitors induces a shift from proliferative to asymmetric neurogenic divisions independently of any CDK interaction but we find that this interaction is required to stimulate neurogenic symmetric terminal divisions (Figure 8A). Our results suggest a dual machinery downstream of CDC25B during the course of neurogenesis (Figure 8B). In one instance CDC25B activity on symmetric neurogenic division is dependent on its interaction with CDK1, while asymmetric neurogenic divisions are promoted by CDC25B independently of its interaction with CDK1, indicating that it involves a new substrate of the phosphatase (Figure 8B).

**Figure 8.**
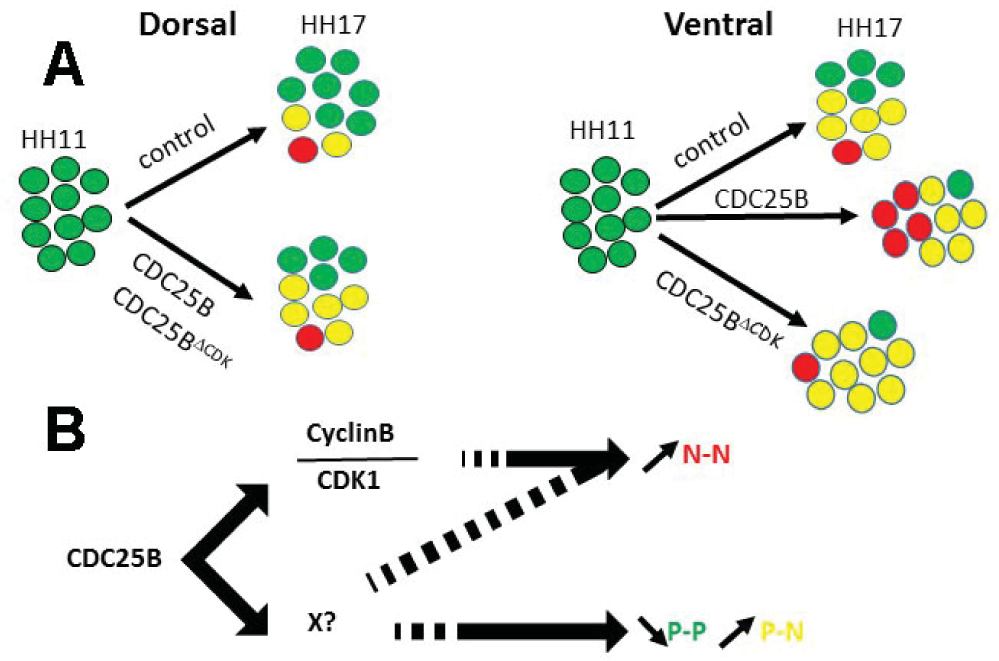
Schematic of CDC25B modes of action. **A**: Different activities of CDC25B on neuroepithelial progenitors. HH11 dorsal neural tube electroporated with control vector, exhibit at HH17 mainly proliferative progenitors schematized using 7 green (PP), 2 yellow (PN) and 1 red ball (NN). CDC25B or CDC25B^∆CDK^ gain-of-function increase asymmetric neurogenic progeny (PN, yellow). In the ventral neural tube, control conditions, lead to a majority of asymmetric neurogenic divisions PN (yellow). CDC25B gain-of-function increases symmetric terminal neurogenic divisions NN (red) whereas CDC25B^∆CDK^ gain-of-function increases asymmetric neurogenic divisions PN (yellow)**. B**: CDC25B dual activity on CDK/cyclinB complexes and/or on an unknown factor X reduces PP progeny and promotes PN progeny or stimulates NN progeny.

### CDC25B is required for efficient neuron production in mammals

In mammals three CDC25s (A, B, C) have been characterized, whereas only two CDC25S (A and B) have been found in chicken (Agius et al., 2015). In mouse, CDC25A loss-of-function is embryonic lethal, whereas loss-of-function of CDC25B or C or both has no apparent phenotype except female sterility (Boutros et al., 2007). Crossing our floxed mice to ubiquitous Cre: PGK-Cre^m^(Lallemand, Luria, Haffner-Krausz, & Lonai, 1998) also results in female sterility (data not shown). CDC25A has been described playing a major role in the G1-S transition and is capable of compensating the loss-of-function of the other CDC25 members. In the mouse embryonic neural tube, both CDC25A and CDC25C display a broad expression pattern, while CDC25B is mainly expressed in domains where neurogenesis occurs (Agius et al., 2015) and Figure 1. The conditional loss-of-function in the mouse CNS, shows for the first time that CDC25B is involved simultaneously in the control of G2 phase length and of spinal neurogenesis. This observation substantiates our data showing that CDC25B downregulation, performed using RNAi in chicken embryo, induces a reduction in neurogenesis (Peco et al., 2012). Two other studies link CDC25B and neurogenesis. First in *Xenopus,* FoxM1 and CDC25B loss-of-function has been shown to reduce expression of neuronal differentiation markers, but not early neuroectoderm markers (Ueno et al., 2008). In this context, epistasic analysis shows that FoxM1 loss-of-function can be rescued by CDC25B gain-of-function (Ueno et al., 2008). Second, MCPH1 knock out mice display a microcephalic phenotype due to an alteration of the Chk1-Cdc25-Cdk1 pathway. Indeed, MCPH1 mutants display a decreased level of the inhibitory Chk1 kinase localized to centrosomes, leading to increased Cdc25B and Cdk1 activities. A premature activation of Cdk1 leads to an asynchrony between mitotic entry and centrosome cycle. This disturbs mitotic spindle alignment, promoting oblique orientation and precocious neurogenic asymmetric divisions (Gruber et al., 2011). Moreover, the reduced neurogenic production in the MCPH1 loss-of-function can be restored by a concomitant Cdc25B loss-of-function, demonstrating the phosphatase's pivotal role in the neurogenic phenotype. Altogether, these observations indicate that CDC25B regulation is broadly used during nervous system development among vertebrate species.

### CDC25B changes the mode of division depending on neural progenitor status

CDC25B downregulation reduces the transition from proliferative to both asymmetric and terminal symmetric neurogenic divisions. To be able to clarify the role of CDC25B on both types of division, we use the cell cycle cis-regulatory element combined with the rapid degradation of CDC25B at the end of the M phase, to reproduce the endogenous cyclic expression of the phosphatase (Korner et al., 2001). In addition, we take advantage of the fact that the progenitor population in the dorsal spinal cord is usually considered younger than its ventral counterpart (Kicheva et al., 2014; Saade et al., 2013), and that neuron production progresses from ventral to dorsal in the neural tube (Peco et al., 2012). Using this paradigm, we show that CDC25B gain-of-function promotes asymmetric or symmetric neurogenic divisions, depending on the population of progenitors targeted. In the dorsal neural tube, CDC25B gain-of-function increases asymmetric neurogenic divisions compared to control conditions, i.e., the phosphatase stimulates the shift from PP to PN divisions (Figure 8A). In the ventral neural tube, the gain-of-function leads to an increase in NN divisions at the expense of PP divisions, the percentage of PN divisions being unchanged (Figure 8A). Based on the quantitative analysis of the progenitor populations in our different conditions, we propose that ectopic expression of the phosphatase can be interpreted in different ways depending on the context, and that the phosphatase's phenotype can be generated in more than one manner. We find that CDC25B has the capacity to convert PP into PN in a young tissue, while in an older tissue CDC25B can convert PP into either PN or NN. With respect to what occurs in an older tissue, either the phosphatase converts PP into PN or NN, or the phosphatase initially promotes PP into PN and subsequently, using the principle of communicating vessels in an older tissue, promotes PN into NN. We speculate that CDC25B acts as a maturating factor in the progression from stem pool to differentiated neurons, and we suggest that this element of the cell cycle machinery has been coopted to regulate independently cell cycle progression and neurogenesis.

### Mathematical modelling of the neuronal fraction in the dorsal neural tube

The model predicts with accuracy the ratio of neuron in the three conditions in the ventral neural tube and in the control condition in the dorsal neural tube. In the latter, in CDC25B and CDC25B^∆CDK^ in-of-functions, the model calculates a larger fraction of neurons than what is observed experimentally (Figure 6C). We have several hypotheses to explain this discrepancy between the predictions and the data. First of all, at HH11, endogenous CDC25B is expressed in the ventral neural tube but not in the dorsal neural tube. This means that electroporation causes a true gain-of-function in the dorsal domain, while in the ventral domain there is only a dosage modification of a component already present. Then, CDC25B regulation is complex, and an active degradation mechanism in the dorsal neural tube could attenuate the gain-of-function. Another possibility is that electroporated gain-of-function, which is also cell cycle dependent, could be less efficient with time and thereby lead to fewer neurons than expected. Alternatively, the signaling pathway downstream of CDC25B could be expressed differently in the ventral and dorsal neural tubes, and this could limit the gain-of-function effect in the dorsal neural tube. All things considered, we regard the discrepancy between our predictions and our data as a challenging milestone that deserves further investigation. One can always formalize an “ad hoc" model for each hypothesis mentioned above in order to fit the observed fractions of neurons, since free parameters can always be adjusted at will. However, we prefer to stress that the standard model for these dynamics still requires identifying other elements in order to reconcile the predictions with the data of this study.

### CDC25B promotes asymmetric neurogenic divisions independently of CDK interaction but symmetric neurogenic divisions require CDK interaction

CDC25B^∆CDK^ turns proliferative divisions into asymmetric neurogenic divisions, but this mutated protein cannot promote symmetric neurogenic divisions (Figure 8A). This result suggests that CDC25B phosphatase affects neurogenesis via two molecular pathways, one dependant and one independent of CDK interaction (Figure 8B). A follow-up to this work could be to characterise the players downstream of CDC25B that are CDK independent. Other CDC25B substrates have been characterised, such as steroid receptors (Ma, Liu, Ngan, & Tsai, 2001), or the peri-centriolar material component Kizuna (Thomas et al., 2014). A recent analysis using microarrayed Tyr(P) peptides representing confirmed and theoretical phosphorylation motifs from the cellular proteome, identifies more than 130 potential CDC25B substrates (Zhao et al., 2015). These substrates are implicated in signalling pathways like Delta/Notch or Wnt, in microtubule dynamics, transcription, epigenetic modifications, mitotic spindle or proteasome activity (Zhao et al., 2015), and all of them could play role in cell fate choice (Akhtar et al., 2009; Aubert, Dunstan, Chambers, & Smith, 2002; Das & Storey, 2012; Gotz & Huttner, 2005; Hammerle & Tejedor, 2007; Jiang & Hsieh, 2014; Kimura, Miki, & Nakanishi, 2014; Li et al., 2012; MuhChyi, Juliandi, Matsuda, & Nakashima, 2013; Olivera-Martinez et al., 2014; Sato, Meijer, Skaltsounis, Greengard, & Brivanlou, 2004; Schwartz & Pirrotta, 2007; Vilas-Boas, Fior, Swedlow, Storey, & Henrique, 2011).

Understanding CDC25B function also depends upon identifying the intracellular localisation of CDC25B activity required for neurogenesis. CDC25B is present in the cytoplasm and/or nucleus according to the cell cycle phase. Moreover, CDC25B protein has been shown to accumulate asymmetrically around the mother centrosome during S and early G2 and it is finally evenly distributed on both centrosomes at late G2 and during mitosis (Boutros & Ducommun, 2008; Dutertre et al., 2004). In mouse Mcph1- deficiency, neurogenesis impairment has been linked to premature activation of Cdc25B expression on centrosomes, leading to imbalanced centrosome maturation and defects in mitotic spindle misalignment (Gruber et al., 2011). As shown here, we could not detect any variation in orientation or size of the mitotic spindle following CDC25B ectopic expression. This suggests that CDC25B centrosomal expression might regulate molecular cascades involved in neurogenesis in parallel or downstream of the mitotic spindle. In this line, Shh induced symmetric recruitment of PKA to the centrosome during neural progenitor divisions, has been involved in promoting expansion of the progenitor pool (Saade et al., 2017). Similarly, Mib1, a known regulator of Notch signalling, has been characterized as an intrinsic fate determinant whose asymmetric localization with centriolar satellite material of proliferating progenitors induces neurogenesis (Tozer, Baek, Fischer, Goiame, & Morin, 2017).

Modifying signalling pathways controlling neurogenesis could also explain the role of CDC25B in promoting symmetric neurogenic divisions that require the interaction between CDC25B and CDK and/or a modification of the G2 phase length. Various experiments have linked G2 phase length with the modulation of signalling pathways such as Wnt or Delta/Notch (Cisneros, Latasa, Garcia-Flores, & Frade, 2008; Davidson et al., 2009; Latasa, Cisneros, & Frade, 2009; Lee, White, Hurov, Stappenbeck, & Piwnica-Worms, 2009; Vilas-Boas et al., 2011). In mouse, CDC25A, B and C triple KO (TKO) exhibits epithelial cells in the small intestine blocked in G1 or G2, accompanied by an enhanced Wnt signalling activity (Lee et al., 2009). Similarly, in Drosophila, the knockdown of String (a CDC25 ortholog in drosophila) results in G2/M arrest and enhances Wnt signalling (Davidson et al., 2009). In neuroepithelial cells, activation of the Notch signalling pathway is regulated by cell cycle progression (Cisneros et al., 2008; Murciano, Zamora, Lopez-Sanchez, & Frade, 2002; Vilas-Boas et al., 2011). Further experiments will be necessary to understand the possible links between CDC25B and the signalling pathways that modulate cell fate decisions during neurogenesis.

In conclusion, we propose that our data illustrate that cell cycle core regulators might have been coopted to elicit additional functions in parallel to cell cycle control. We show that a positive cell cycle regulator, CDC25B, unexpectedly promotes differentiation and reduces proliferative divisions. Cell cycle regulators are routinely described as deregulated in cancers and are associated with increased proliferation. Understanding their function outside the cell cycle is therefore crucial to characterising their molecular and cellular mechanisms of action and to foresee novel therapeutic strategies.

## MATERIALS AND METHODS

### Embryos

Fertile chicken eggs at 38°C in a humidified incubator yielded appropriately staged embryos (Hamburger & Hamilton, 1992). Animal related procedures were performed according to EC guidelines (86/609/CEE), French Decree no. 97/748 and the CNRS recommendations.

### Generating a *Cdc25B* floxed allele and a *CDC25B*^*nesKO*^ littermates

Experiments were performed in accordance with European Community guidelines regarding care and use of animals, agreement from the Ministère de I'Enseignement Supérieur et de la Recherche number: C3155511, reference 01024.01 and the CNRS recommendations. To disrupt *Cdc25B* function, we generated a modified allele of *Cdc25B* (Mouse Clinical Institute, IGBMC, lllkirch). Using Homologous recombination in embryonic cells (ES), we inserted two LoxP sites, flanking exon 4 to exon 7 of the *Cdc25B* gene (referred to as Floxed allele). Upon Cre-mediated excision exons 4 to 7 are deleted and following intron splicing, a premature stop codon is generated, leading to a truncated protein of 134 aa. The activity of this remaining peptide has been tested in a cellular model and has no activity (not shown). We first generated a mutant mouse line **(Cdc25B^-)** by crossing *Cdc25B* floxed mice with *PGK-Cre* mice, resulting in an ubiquitous and permanent deletion of *Cdc25B.* In order to delete Cdc25B activity specifically at the onset of neurogenesis, we crossed Cdc25B^fl/-^ mice with transgenic mice expressing the Cre recombinase under the control of the rat Nestin (Nes) promoter and enhancer (Tronche et al., 1999). The effect of expressing Cre recombinase on proliferation and neurogenesis was evaluated by comparing *Cdc25B*^*fl/+*^ and *NestinCre;Cdc25B*^*fl/+*^ littermates. As there were no phenotypic differences between these embryos for any of the parameters that we measured (not shown), they were both included with the *Cdc25B*^*fl/-*^ littermates in the control group.

### Statistical analysis of the mouse neuronal phenotype

For each experiment, at least three independent litters and three different slides per embryo were analyzed. To compare the number of neuron between control and conditional mutant embryos, we used a statistical model called the “mixed effect model”. This model contains both the fixed effect i.e., the genotype of the embryo (control or conditional mutant) - and random effects i.e., the variability induced by the age of the litter and by the embryo nested in the litter. Random effects were excluded using the R software and the package “nlme”, and we applied the following formula:

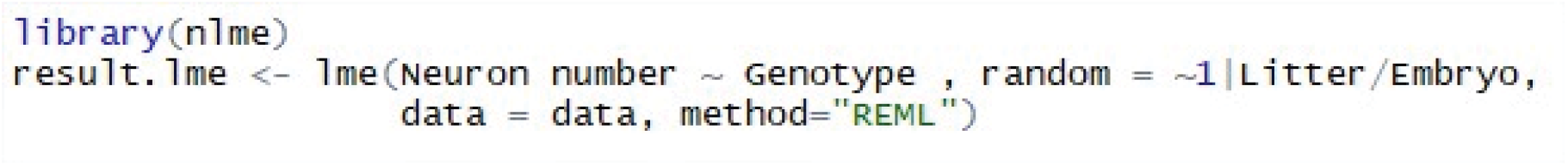

To test the effect of the genotype on the number of neuron, we next performed an ANOVA test. *p < 0.05; ** p < 0.01; *** p < 0.001

### DNA constructs and *in ovo* electroporation

*In ovo* electroporation experiments were performed using 1.5- to 2-day-old chickens as described previously (Peco et al., 2012). Loss and gain of function experiments were performed using a vector expressing the various human CDC25 isoforms (hCDC25B3, hCDC25B3^∆CDK^ hCDC25B3^∆P∆CDK^) under the control of a cis regulatory element of the mouse Cdc25B called pccRE. A control vector was generated with the βGal gene downstream of the pccRE. All gain-of-function experiments were performed at 1.5 μg/μl. The Sox2p-GFP, Tis21p-RFP, and NeuroD-luciferase constructs were obtained from E. Marti and used at 1 μg/μl, 0.5 μg/μl and 1 μg/μl, respectively.

### *In situ* hybridization and immunohistochemistry on mouse and chick embryos

Mouse embryos were dissected in cold PBS and fixed in 4% paraformaldehyde overnight at 4°C. Then they were embedded in 5% low melting agarose before sectioning on a Leica vibratome, in 50 μm thick transversal sections. *In situ* hybridization was performed as published (Lacomme, Liaubet, Pituello, & Bel-Vialar, 2012). Riboprobes to detect mCdc25B transcripts were synthesized from linearized plasmid containing the full Cdc25B cDNA (riboprobe sequence available on request). Immunohistochemistry was performed as described in (Lobjois, Benazeraf, Bertrand, Medevielle, & Pituello, 2004). The antibodies used were the anti-Pax2 (Covance), guinea pig anti-Tlx3 (gift from C.Birchmeier (Muller et al., 2005)) and anti-Pax7 (Hybridoma Bank). For chick embryos, proteins or transcripts were detected on 40 μm vibratome sections, as previously described (Peco et al., 2012). The antibodies used were: anti-HuC/D (Molecular Probes), anti-Sox2 (Chemicon), anti-PH3 (Upstate Biotechnology), anti-BrdU (mouse monoclonal, G3G4), anti-BrdU (rat anti-BrdU, AbD Serotec), anti-active caspase 3 (BD Biosciences), and anti-GFP (Invitrogen).

### Cell proliferation and survival analyses

Cell proliferation was evaluated by incorporation of 5-ethynyl-2’-deoxyuridine (Click-iT EdU Alexa Fluor 647 Imaging Kit, Invitrogen). 10 μl of 250 μM EdU solution were injected into chicken embryos harvested 30 minutes later, fixed for one hour and processed for vibratome sectioning. EdU immunodetection was performed according to manufacturer's instructions. Mitotic cells were detected using anti-PH3. G2-phase length was determined using the percentage of labeled mitoses (PLM) paradigm (Quastler & Sherman, 1959). EdU incorporation was performed as described above, except that a similar dose of EdU was added every 2 hours, and embryos were harvested from 30 to 180 minutes later. Embryos were fixed and labeled for both EdU and PH3. We then quantified the percentage of PH3 and EdU co-labeled nuclei with increasing times of exposure to EdU. The progression of this percentage is proportional to G2-phase duration. Cell death was analyzed by immunofluorescence, using the anti-active Caspase 3 monoclonal antibody (BD Biosciences).

### EdU incorporation in mice

For EdU staining experiments in mouse, 100 μl of 10mg/ml EdU were injected intraperitoneally into pregnant mice. Litters were harvested 1, 2 or 3 hours following injection.

### Imaging and data analysis

Slices (40 μm) were analyzed using a SP5 Leica confocal microscope as described previously (Peco et al., 2012). Experiments were performed in triplicate. For each embryo, confocal analyses were performed on at least three slices. Confocal images were acquired throughout the slices at 3 μm *z* intervals.

### Tis21::RFP/Sox2::GFP Quantification

For each experimental slice, Z sections were acquired every 3 μm, and blind cell quantifications were performed on one out of every three Z sections to avoid counting the same cell twice. For each slice, the percentage of PP, PN and NN divisions is determined using the sum of counted Z sections. For each experimental condition, the number of embryos analyzed and of cells counted is indicated in the Figure legend.

### In Vivo Luciferase Reporter Assay

Embryos were electroporated with the DMAs indicated together with a NeuroDp-Luciferase reporter (Saade et al., 2013) and with a renilia-construct (Promega) for normalization. GFP-positive neural tubes were dissected out at 48 hours after electroporation and homogenized in passive lysis buffer. Firefly- and renilla-luciferase activities were measured by the Dual Luciferase Reporter Assay System (Promega), and the data are represented as the mean ± sem from at least 14 embryos per experimental condition.

### Statistics

Quantitative data are expressed as mean ± s.e.m. Statistical analysis was performed using the GraphPad Prism software. Significance was assessed by performing ANOVA followed by the Student- Mann-Whitney test, (\**P*<0.05, \*\**P*<0.01, \*\*\**P*<0.001, ****P<0.0001 and n s non significant).

## ACKNOWLEDGMENTS

We are grateful to Dr. Elisa Marti for sharing plasmids. We thank Drs. Bertrand Bénazéraf, Alice Davy, Bernard Ducommun, Xavier Morin and Alain Vincent for critical reading of the manuscript and Dr. Caroline Monod for improving the English. We thank the CBI animal facilities and the Toulouse Regional Imaging platform (TRI) for technical support. We acknowledge the Developmental Studies Hybridoma Bank, created by the NICHD of the NIH and maintained at The University of Iowa, Department of Biology, Iowa City, IA 52242 for supplying monoclonal antibodies.

## Supplemental Information Text 1^1^ — Modeling the dynamics

Azais Manon, Gautrais Jacques*

*

### Abstract

We present the model of the dynamics for the interpretation of CDC25B experiments.

We present the solution when fate parameters are considered steady over the time window of the analyses.

We present the sensitivity of the dynamics to the modes of division.

We present and explain the predicted fractions of neurons under the three conditions and the two zones.

^1^for the paper: NEUROGENIC DECISIONS REQUIRE A CELL CYCLE INDEPENDENT FUNCTION OF THE CDC25B PHOSPHATASE, Fr´ed´eric BONNET, M´elanie ROUSSAT, Angie MOLINA, Manon AZAIS, Sophie BEL-VIALAR, Jacques GAUTRAIS, Fabienne PITUELLO and Eric AGIUS.

### 1 The model

We consider a population of cells *C*(*t*) at time *t,* part of which are proliferating progenitors *P*(*t*), part of which are differentiated neurons *N*(*t*), with

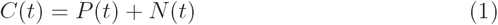

The dividing progenitors can undergo three kinds of fate, yielding:

- some proliferative divisions ending with two progenitors (pp-divisions)
- some asymmetric divisions ending with one progenitor and one neuron (pn-divisions)
- some terminal divisions ending with two neurons (nn-divisions)

We consider that the division of a cell in two cells is instantaneous (it is always possible to find a date before which there is one cell, and after which there are two cells).

We also consider that division events occur uniformly in time (asynchronously).

Let us denote :

*η* the rate at which P-cells undergo divisions (in fraction of the P-pool per unit time)

*α*_pp_(*t*) the fraction of dividing cells undergoing pp-divisions

*α*_pn_(*t*) the fraction of dividing cells undergoing pn-divisions

*α*_nn_(*t*) the fraction of dividing cells undergoing nn-divisions

*P*(0)*, N*(0) the quantity of P-cells and N-cells known at time *t* = 0.

In general, the fractions of pp-, pn- and nn-divisions can evolve with time, under the constraint that *α*_*pp*_ + *α*_*pn*_ + *α*_*nn*_ = 1, and so might as well the division rate.

The time change 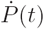 of pool *P(t)* (resp. 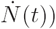 is then driven at time *t* by:

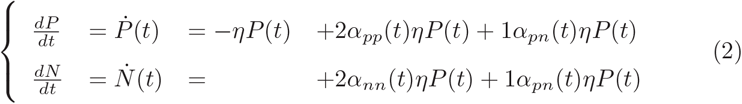

where in the first equation :

- *-ηP*(*t*) quantifies the rate at which P-cells disappear from the pool *P(t)* because they divide. The quantity of disappearing P-cells between *t* and *t + dt* is then *ηP(t)dt.*
- *α*_*pp*_*ηP*(*t*) quantifies the fraction of this quantity that undergoes a pp-division; it doubles to yield 2 P and adds up to the pool P(t) (hence the factor 2)
- *α*_*pn*_*ηP*(*t*) quantifies the fraction of this quantity that undergoes a pndivision; it doubles to yield 1 P and 1 N, so only half (the P part) adds up to the pool P(t) (hence the factor 1)

correspondingly in the second equation :

- *α*_*nn*_*ηP*(*t*) quantifies the fraction of this quantity that undergoes a nn-division; it doubles to yield 2 N and adds up to the pool N(t) (hence the factor 2)
- *α*_*pn*_*ηP*(*t*) is the fraction of this quantity that undergoes a pn-division; it doubles to yield 1 P and 1 N and only half (the N part) adds up to the pool N(t) (hence the factor 1)

### 2 Solutions with unvarying parameters

Considering a period of time during which the fractions of pp-, pn- and nn-divisions do not evolve with time, the dynamics can be written:

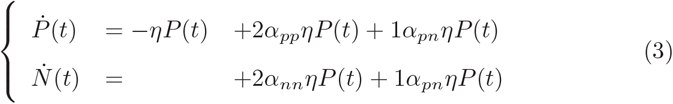

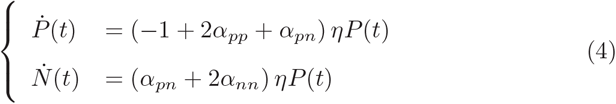

Let *γ* = −1 + 2*α*_*pp*_ + *α*_*pn*_.

Considering that *α*_*pp*_ + *α*_*pn*_ + *α*_*nn*_ = 1, we have:

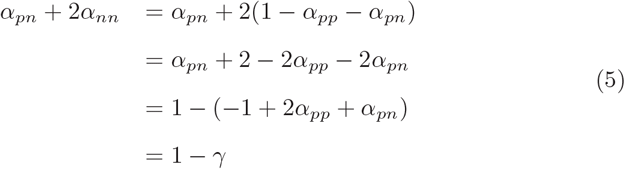

Hence,

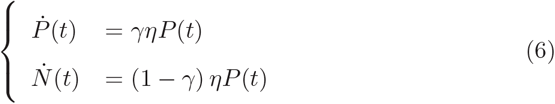

and the solutions are of the general form:

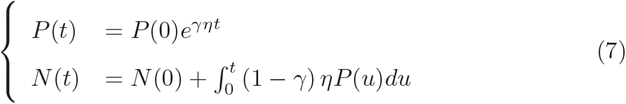

plugging the first into the second, we have:

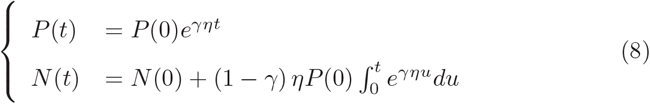

#### 2.1 Explicit solutions

For explicit solutions, we have to consider two cases: *γ* = 0 and *γ* ≠ 0.

Fo r *γ* = 0, we have:

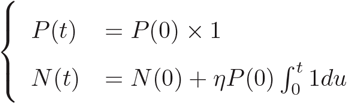

so that:

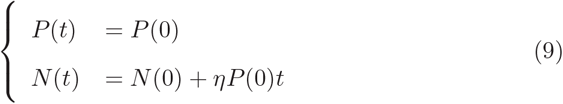

In that case, the pool of progenitors is steady, and the pool of neurons increases linearly with time.

Fo r *γ* = 0, solving the integral in the second equation yields:

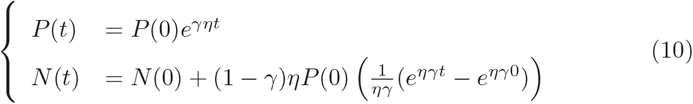

so that:

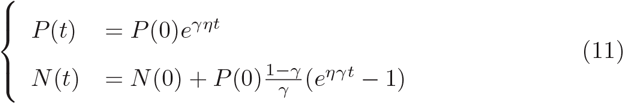

In that case, the evolution of the system depends on the sign of *γ*.

#### 2.2 Meaning of γ

We note that, for a given mitosis rate *η*, the dynamics only depend upon *γ*.

We have *γ* = 2*α*_*pp*_ + *α*_*pn*_ — 1 = 2*α*_*pp*_ + *α*_*pn*_ — (*α*_*pp*_ + *α*_*pn*_ + *α*_*nn*_) = *α*_*pp*_ — *α*_*nn*_.

The case *γ* = 0 (Eq.9) corresponds to *α*_*pp*_ = *α*_*nn*_. Here, the P-pool is steady and can be considered as a source of N-cells emitted at the steady rate *ηP*(0) (N-cells per unit time):

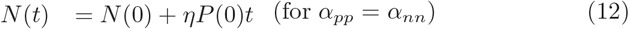

The case *α*_*pp*_ > *α*_*nn*_ yields *γ* > 0, so that the P-pool will increase with time. At the extreme, a purely proliferative P-pool corresponds to *α*_*pp*_ = 1 and *αnn* = 0, hence *γ*=1. In that case, the dynamics simplify to the classical proliferative equation for the P-pool, while the N-pool remains unchanged:

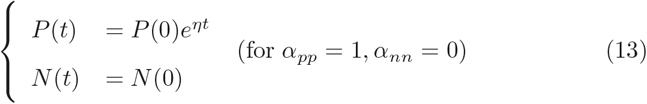

The case *α*_*pp*_ < *α*_*nn*_ yields *γ* < 0, so that the P-pool will decrease with time. At the extreme, a fully differentiating P-pool corresponds to *α*_*pp*_ = 0 and *α*_*nn*_ = 1, hence *γ* = — 1. In that case, the P-pool undergoes a classical exponential decay, and the N-pool increases in proportion of the remaining P-pool, up to 2*P*(0):

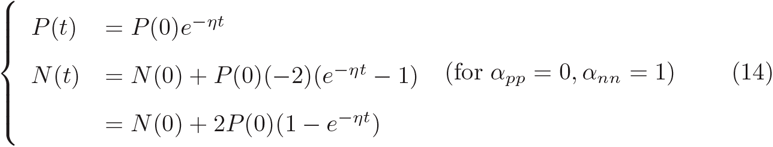

Regarding the total population *C*(*t*) = *P*(*t*)+*N*(*t*) (fig. 1), positive (or null) value of *γ* (*α*_*pp*_ ≥ *α*_*nn*_) allows an infinite growth of the total population *C*(*t*) whereas the growth saturates as soon as *γ* < 0 (*α*_*pp*_ < *α*_*nn*_). We note here that we made the hypothesis that the fate parameters were considered as steady over time, so interpretations for the real biological system should take into account that these fate parameters actually change over longer time in the real system.

**Figure 1:**
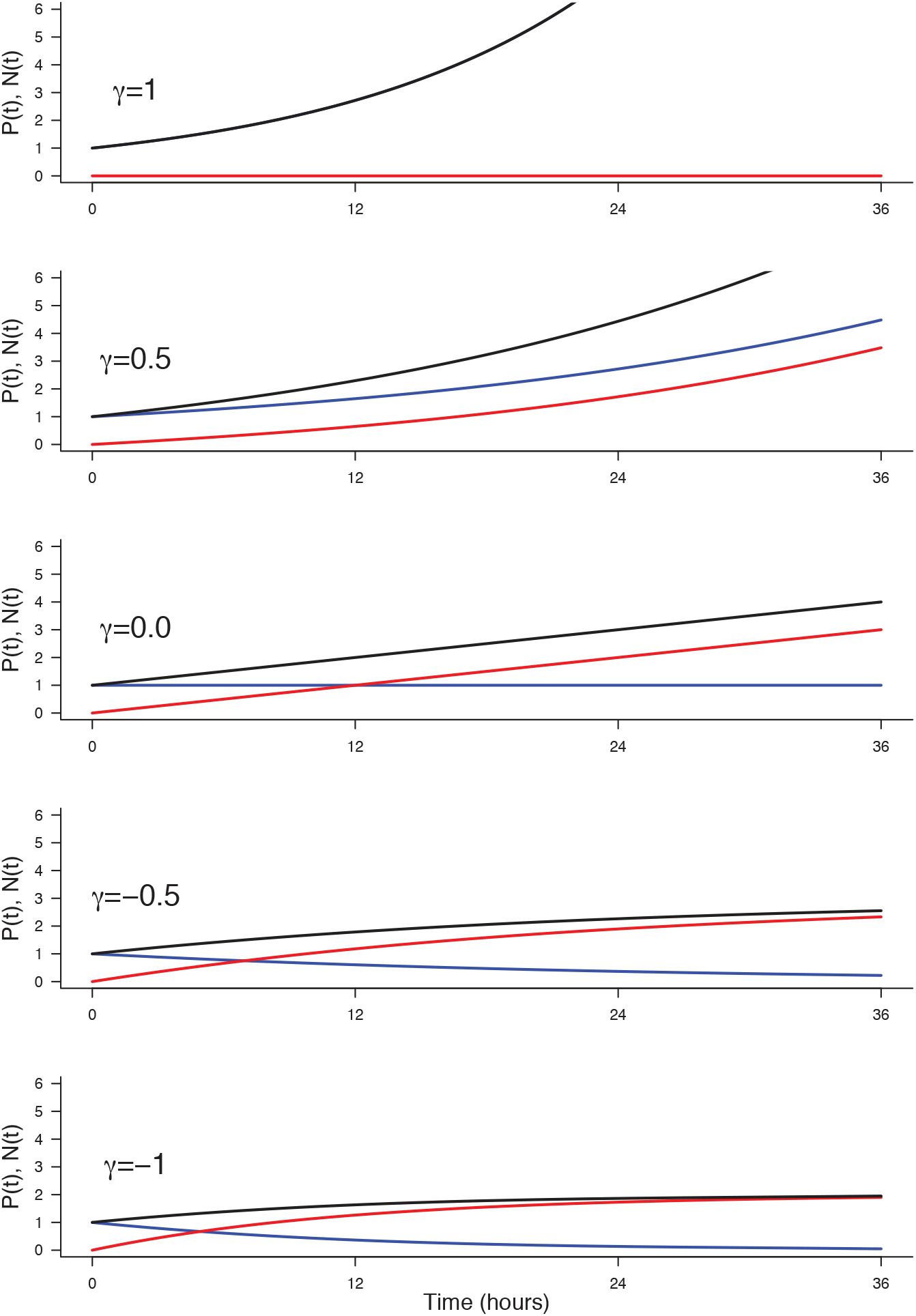
**Effect of *γ* on the evolution of *P*(*t*) (blue), *N*(*t*) (red) and *C*(*t*) = *P(t*) + *N*(*t*) (black).** Parameters used: *P*(0) = 1, *N*(*0*) = 0, *η* = 1*/*12, corresponding to a cycle time of 12 hours.

Regarding the fraction of neurons in the population, *N*(*t*)/*C*(*t*) (fig. 2), it increases as soon as *γ* < 1, yet at a rate depending on *γ*.

**Figure 2:**
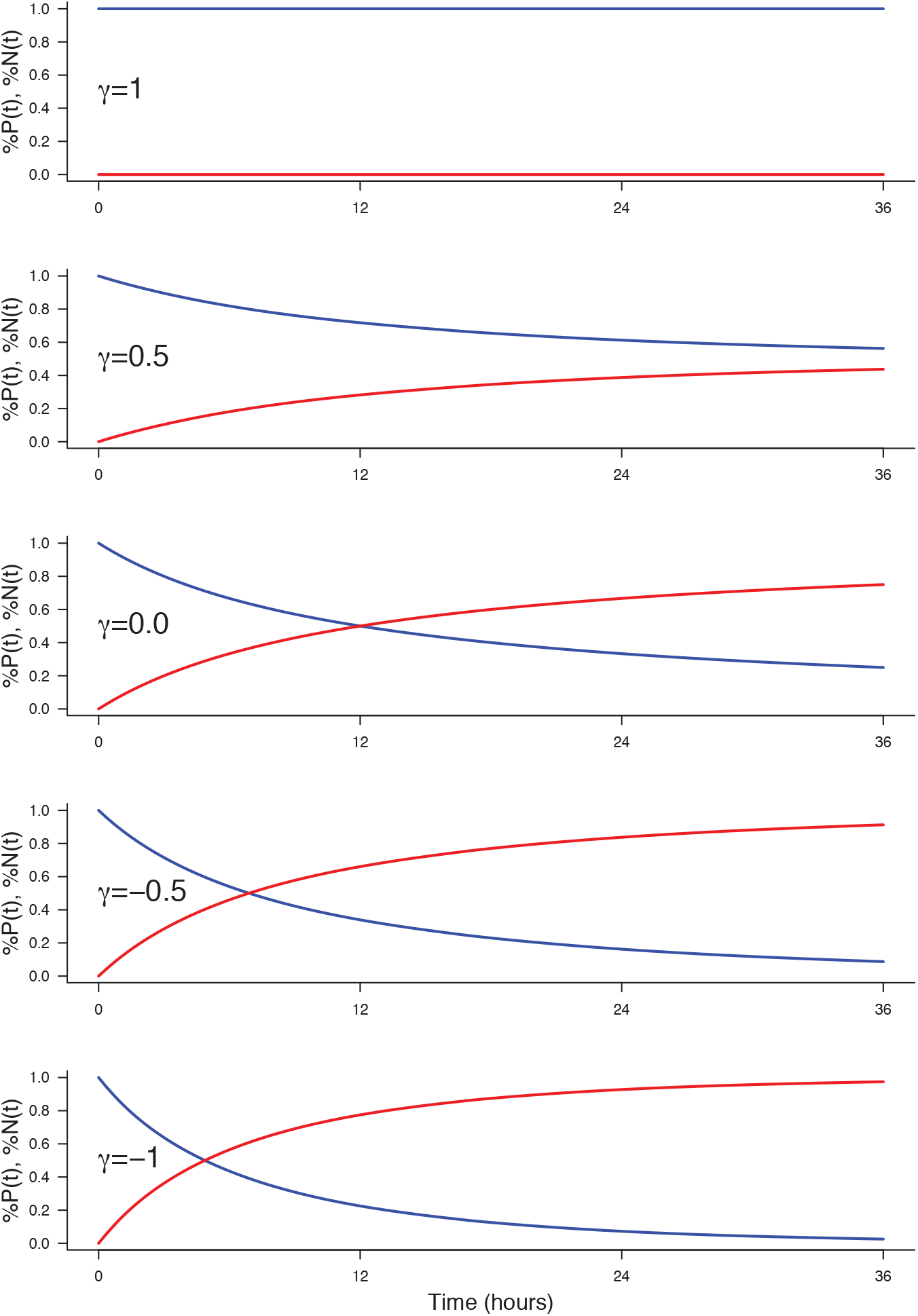
Effect of *γ* on the evolution of the fractions *P*(*t*)/*C*(*t*) (blue) and *N*(*t*)/*C*(*t*) (red).

### 3 Interpretations at the individual cell scale

We have so far describe the system at the population scale. At the individual scale, two different kinds of process (at least) would result in the same dynamics at the population scale described in Eq.2.

#### 3.1 Probabilistic fates, with a common deterministic division rate

The most immediate interpretation is to consider that all cells undergo mitosis at the same rate, and that the fate of any mitosis is stochastic and probabilistically distributed according to (*α*_*pp*_, *α*_*pn*_, *α*_*nn*_). In that case, only the rate *η* (used in the equations at the population scale) has to be determined from cell-scale model, since it depends upon the characteristic time *τ*_*m*_ between two mitosis at the cell scale.

Let us consider the hypothesis that mitosis happen exactly every *τ*_*m*_ for all cells (common deterministic division time), still asynchronously so that division dates are uniformly distributed over time (this is the most common hypothesis in the community). We want to express *η* as a function of *τ*_*m*_.

For the sake of simplicity, let us consider the pure proliferative process (*α*_*pp*_ = 1) so that we deal with only one population *P*(*t*).

Let us start at time 0 with an initial pool *P*_1_(0) containing a very large number of cells (so that *P*_1_(*t*) can be considered as continuous). Since mitosis take a fixed time *τ*_*m*_, their last division occurred before *t =* 0, the oldest division happened at 0–*τ*_*m*_ and they all will make a mitosis in [0 … 0+*τ*_*m*_]. Since divisions are uniformly distributed over time, the number doing a mitosis during a small time interval Δ*t* is proportional to Δ*t*/*τ*_*m*_ and *P*(0). Hence, the loss in *P*_1_ between *t* and *t* + Δ*t* is given by:

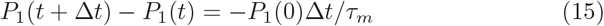

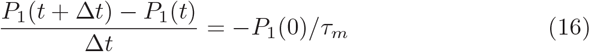

Taking the limit Δ*t →* 0 yields:

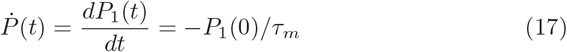

Considering *P*_1_(0), we then have:

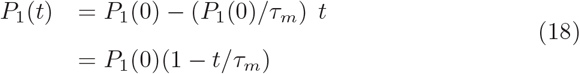

Logically, *P*_1_(*t*) decreases linearly from *P*_1_(0) down to 0 at time *t* = *τ*_*m*_. Meanwhile, the output of each division will populate the next generation, say *P*_2_(*t*), at twice the rate *P*_1_ disappears, up to 2*P*_1_(0) at time *t* = *τ*_*m*_, from which *P*_2_ will start decreasing doing mitosis and populate the third generation *P*_3_ and so on. Such a process would then translate into a population growth which is piecewise linear (fig 3), but very close to an exponential growth. If we equate at time *τ*_*m*_ the piecewise growth, and its exponential approximation at rate *η*, we have:

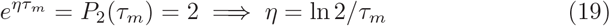

Denoting *τ*_*c*_ = 1*/η* the characteristic time at the population scale, we then have: *τ*_*c*_ = *τ*_*m*_/ln2. Hence, from an observed time *τ*_*c*_ at the population scale, we should infer (under this model) that *τ*_*m*_ = *τ*_*c*_ ln 2, i.e. *τ*_*m*_ ≃ 0.7*τ*_*c*_ (e.g. if population cycle time is 12h, cell cycle time should be around 8h20).

**Figure 3:**
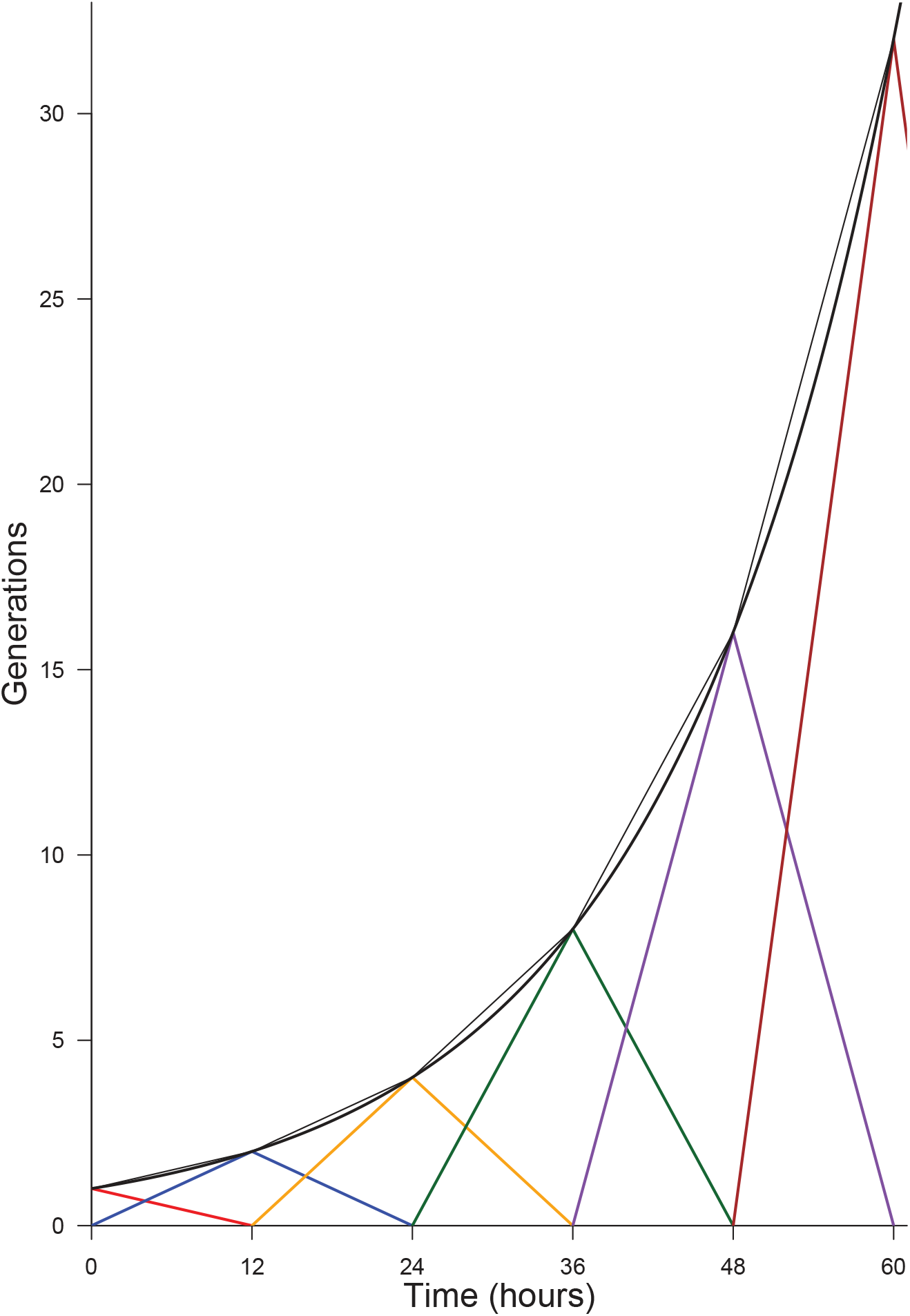
Generations produced by an initial pool *P*_1_(0) = 1, under the hypothesis of a common deterministic division time *τ*_*m*_ = 12 h. Each generation is reported by a color. The thin black curve indicates the total pool present at time *t* (adding the two generations). The thick black curve reports the continuous approximation exp(ln 2 *t*/*τ*_*m*_) (eq. 19)

#### 3.2 Deterministic fates, with specific division rates

Another way to produce the dynamics described in eq.2 at the population scale is to consider that each kind of fate result from a specific division time. In such a picture, the time needed to achieve a cycle deterministically determines the kind of fate.

To exhibit this interpretation, we rewrite eq.2 as follows:

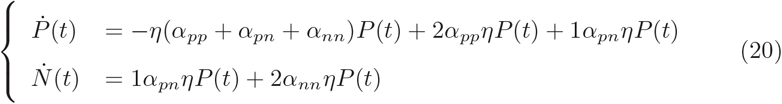

Denoting *η*_*pp*_ = *α*_*pp*_*η* (and correspondingly for *η*_*pn*_ and *η*_*nn*_), we then have:

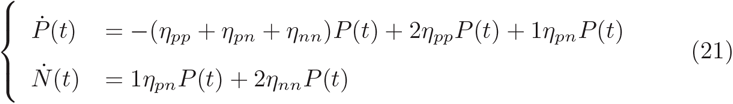

The interpretation is then that, from the pool P(t), the cells leaving it at rate *η*_*pp*_ yield pp-divisions, those leaving it at rate *η*_*pn*_ yield pn-divisions, and the others, leaving it at rate *η*_*nn*_, yield nn-divisions. Overall, the pool *P*(*t*) depletes at the sum rate *η* = *η*_*pp*_ + *η*_*pn*_ + *η*_*nn*_.

Correspondingly, the population cycle time *τ*_*c*_ = 1/*η* would then be given by:

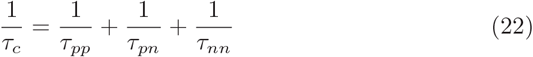

equivalently by:

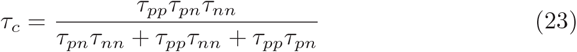

We also note that the distribution of fates is then completely constrained by the *τ*_*pp*_, *τ*_*pn*_, *τ*_*nn*_ (under the constraint that mitosis events are uniformly distributed in time). Indeed, it remains true that the quantity leaving the P-pool during Δ*t* to make pp-divisions is proportional to Δ*t*/*τ*_*pp*_ (corr. for other fates). This implies in turn that the fraction *α*_*pp*_ leaving for an pp-division is *τ*_*c*_/*τ*_*pp*_, correspondingly, *α*_*pn*_ = *τ*_*c*_/*τ*_*pn*_ and *α*_*nn*_ = *τ*_*c*_/*τ*_*nn*_.

As a consequence, if we have experimental measures of *τ*_*c*_ and of a distribution among fates *α*_*pp*_, *α*_*pn*_, *α*_*nn*_, we must conclude that:

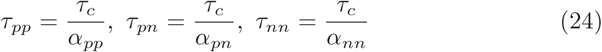

For *τ*_*c*_ = 12 *h,* and a distribution (0.6, 0.3, 0.1), we would obtain:

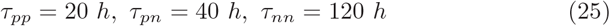

The main point is then: if the ratios between fractions of fate *α*_*pp*_, *α*_*pn*_, *α*_*nn*_ resulted only from differences in rates *η*_*pp*_, *η*_*pn*_, *η*_*nn*_, the ratios between rates must be the same as the ratios between fractions:

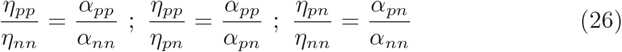

With *α*_*pp*_ = 0.6, *α*_*nn*_ = 0.1, we would have *τ*_*nn*_ = (*α*_*pp*_/*α*_*nn*_)*τ*_*pp*_ = 6 *τ*_*pp*_.

If we exclude the possibility that a nn-division is 6 times as long as a pp-division, then the distribution of fates can not be exclusively determined by differences in fate-based cycle times. It does not exclude that a given kind of fate (e.g. proliferative divisions pp) would require a longer time to be achieved than others, it excludes that such differences would suffice *per se* to explain the differences between the fractions of fates.

### 4 Model predictions using (noisy) data

We obtain experimental measures upon this system at different times after elec-troporation (time 0h): the fractions f_*N*_(24) of neurons at 24h and f_*N*_(48) at 48h (the fraction among the electroporated cells), the distribution of fates at 24h as well as an estimate of *τ*_*c*_ = 12 hours. We make the hypothesis that the fate distribution is steady between 24h and 48h after electroporation, i.e. the 24 hours between the quantification of the mode of division and progenitors and neurons counting. We use the model to check the consistency of these data with the model.

#### 4.1 Knowing the fractions of neurons at 24h and 48h, confidence intervals upon the fate distribution

The first test of consistency was to determine the ranges of distribution of fates which was able to explain the transition from f_*N*_(24) to f_*N*_(48).

If we had a system with only symmetric divisions (e.g. some value for *α*_*pp*_, *αnn* = 1 — *α*_*pp*_, with *α*_*pn*_ = 0), we first ensured that one pair (f_*N*_(24),f_*N*_(48)) would be compatible with only one fate distribution.

Considering *P*(24) + *N*(24) = 1 arbitrary total amount of cells at 24h, we can plug *N*(24) = f_*N*_(24) and *P*(24) = 1 — f_*N*_(24) into eq.11 and get:

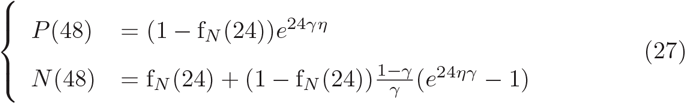

where *P*(48)*, N*(48) correspond to the amount obtained at 48h from this arbitrary amount of 1 at 24h. We have f_*N*_(48) = *N*(48)/(*N*(48) + *P*(48)), yielding :

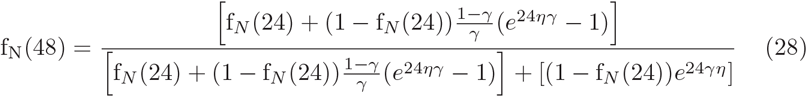

which holds for any initial cell amount (fig. 4).

**Figure 4:**
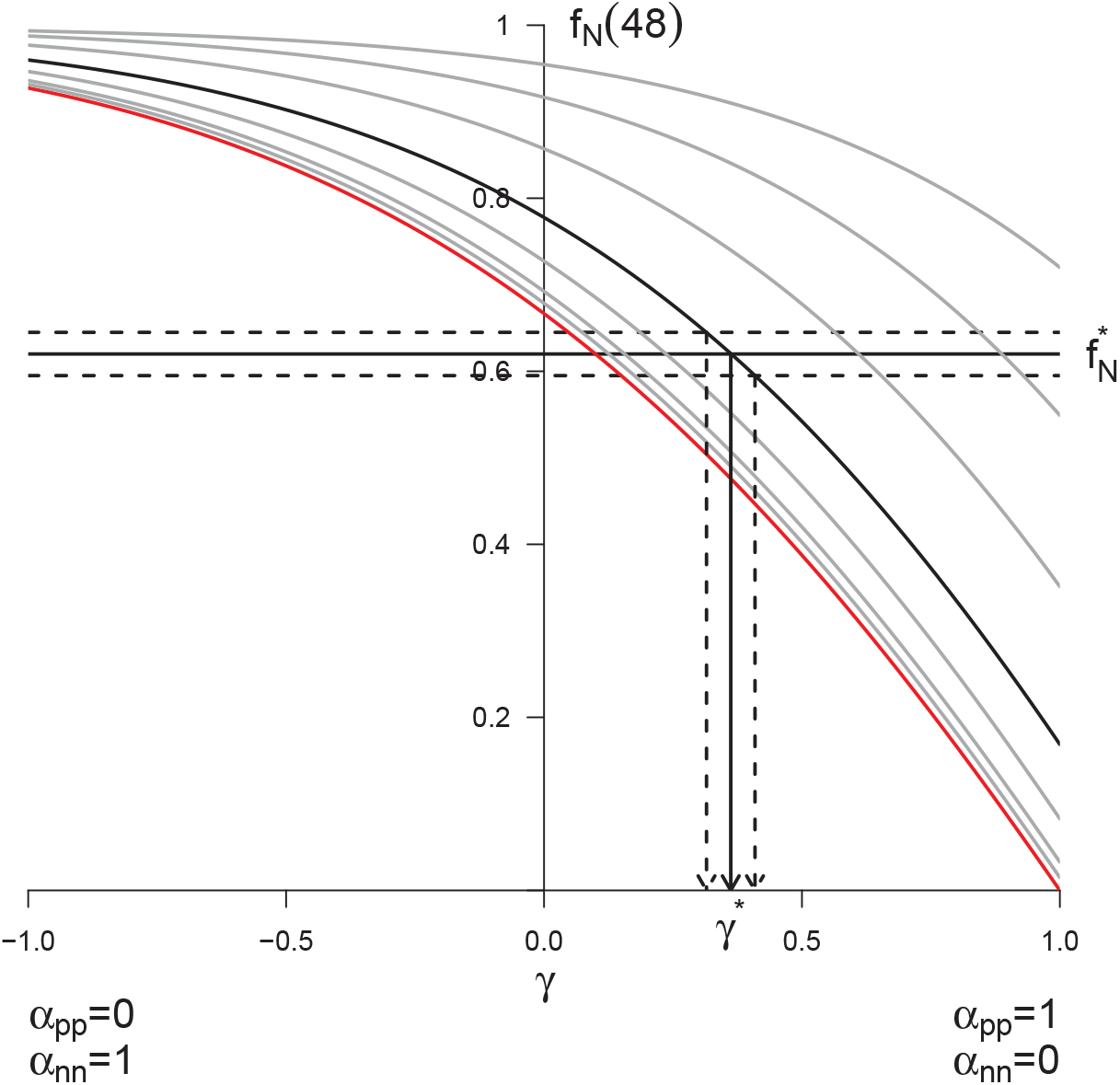
Predicted f_*N*_(48) **from** f_*N*_(24) **for every distribution of symmetric division.** The different curves correspond to different starting values f_*N*_(24) taken in (0.0, 0.1, 0.2, 0.4, 0.6, 0.8, 0.9, 0.95). The bold line corresponds to f_*N*_(24) = 0.6, the red line to f_*N*_(24) = 0.0. Each curve reports the predicted value for f_*N*_(48) starting from the corresponding f_*N*_(24), and for all possible distributions of fates given by *γ* = *α*_*pp*_ - *α*_*nn*_ (x-axis). Each combined (f_*N*_ (24), *γ*) yields only one predicted f_*N*_(48). Conversely, experimental values for the pair (f_*N*_(24), f_*N*_(48)) allow to retrieve the corresponding *γ* theoretical value. As an example, the value corresponding to the arbitrary value 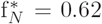 was retrieved numerically using Eq.28. We found *γ*\* = 0.362, yielding *α*_*pp*_ = 0.681 and *α*_*nn*_ = 0.319. Confidence interval upon the distributions of fates can also be drawn using the experimental noise about f_*N*_(48), as illustrated here considering 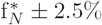.

Now considering the full system with the three kinds of division, there is more than a unique triplet (*a*_*pp*_, *a*_*pn*_, *a*_*pn*_) that are compatible with the unique value of observed (f_*N*_(24), f_*N*_(48)). For instance, less nn-divisions can be compensated for by more pn-divisions, yielding the same *f*_*N*_ (48).

We used the model in the same spirit as in fig.4 to compute the predicted values for f_*N*_(48) for all possible fate triplets. For the system with symmetriconly divisions above, the space of parameters for division is one-dimensional: 7 corresponds to one value of *α*_*pp*_, which constrains in turn the value of *α*_*nn*_. With the three kinds of division, this space of parameters becomes two-dimensional: we need to fix *α*_*pp*_ and *α*_*nn*_, and *α*_*pn*_ is then constrained. Hence the predictions should be drawn over a two-dimensional map.

We compute those maps for each experimental condition, starting from the corresponding observed value f_*N*_(24) (fixing the observed initial condition corresponds here to draw only the bold curve in fig.4). Then, we determine numerically the subset of fate triplets compatible with the 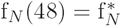 measured in the condition. We also determined numerically the confidence regions for the distributions of fates that can yield 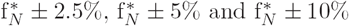.

In the end, we also report the distribution of fates that was actually measured, and check in which confidence interval it is.

**Figure 5:**
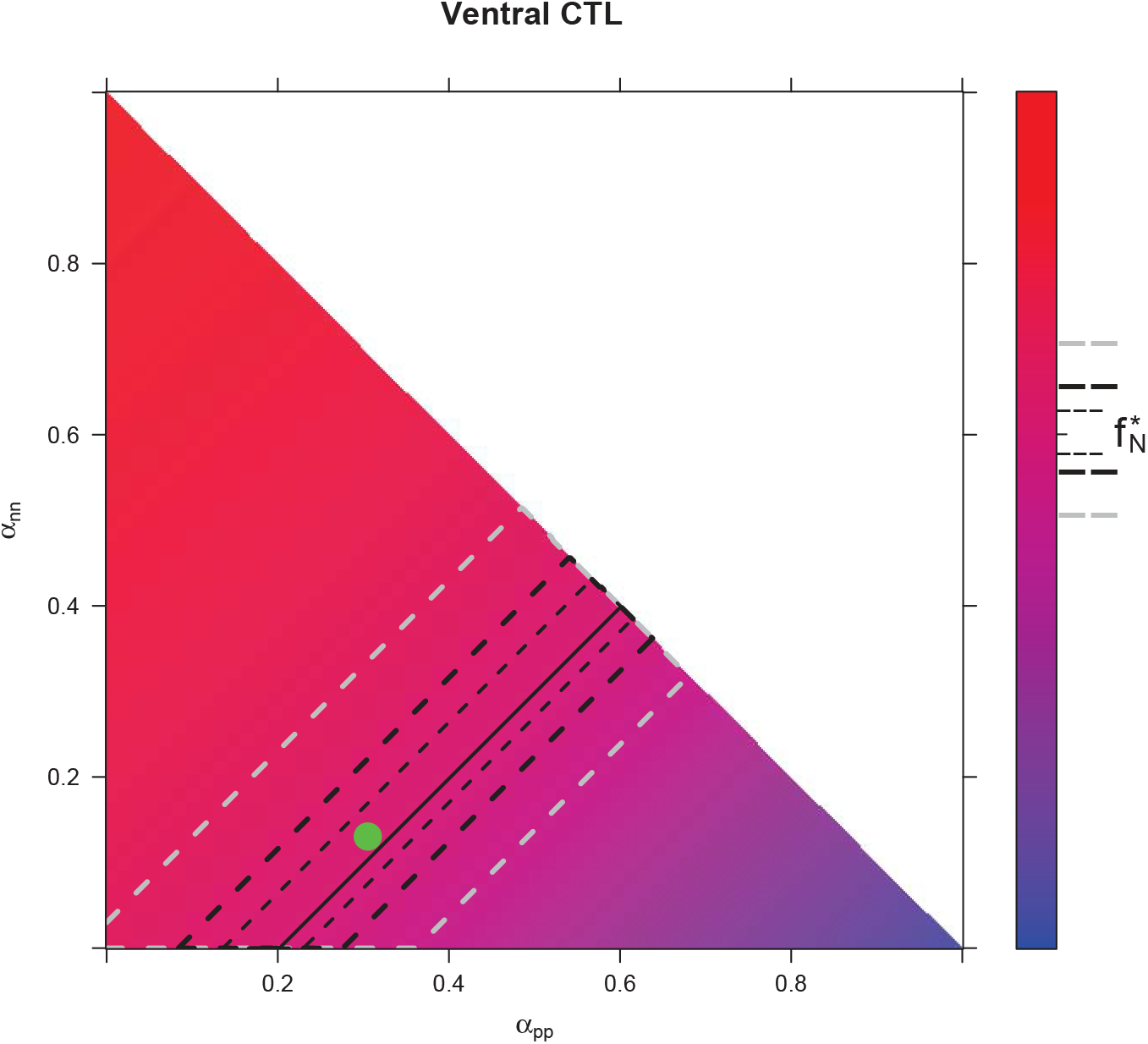
Predicted f_*N*_(48) from f_*N*_(24) for every distribution of fates for control condition in Ventral area. The color scale indicates f_*N*_(48). It is computed from the model, starting from the experimental value of f_*N*_(24) in the prevailing condition, and using all possible distributions of fates *α*_*pp*_ (x-axis), *α*_*nn*_ (y-axis) and *α*_*pn*_ = 1–*α*_*pn*_–*α*_*nn*_. The upper side of the triangle corresponds to *α*_*pn*_ = 0. Confidence interval upon the predicted distributions of fates are drawn for the experimental value 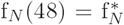. Plain line: all distributions of fates giving exactly 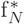. Region delimited by thin dotted line: all distributions of fates compatible with 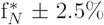, thick dotted line : 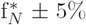, gray dotted line: 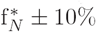. Green dot: observed distribution of fates.

**Figure 6:**
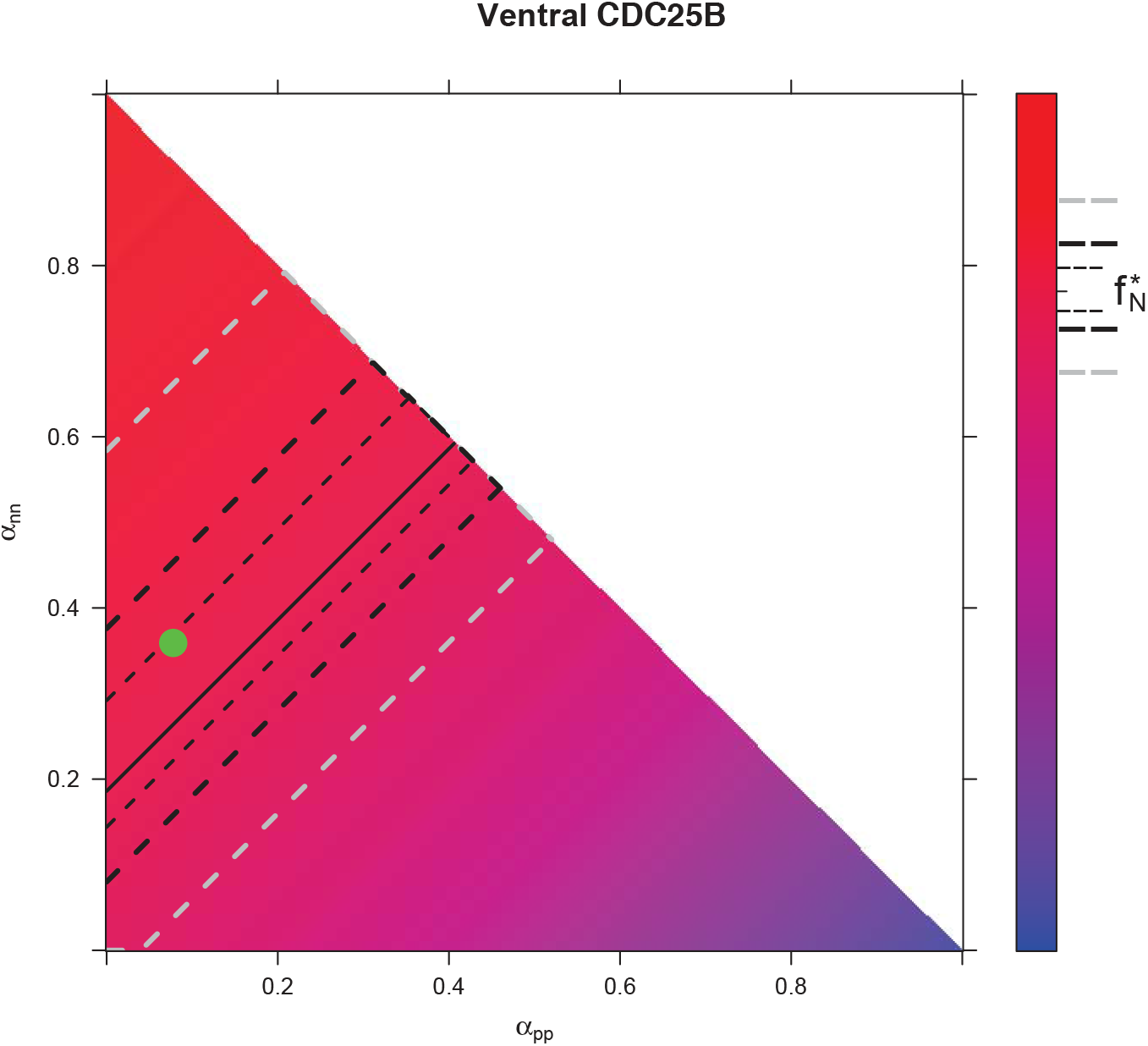
Predicted f_*N*_(48) from f_*N*_(24) for every distribution of fates for CDC25B condition in Ventral area.

**Figure 7:**
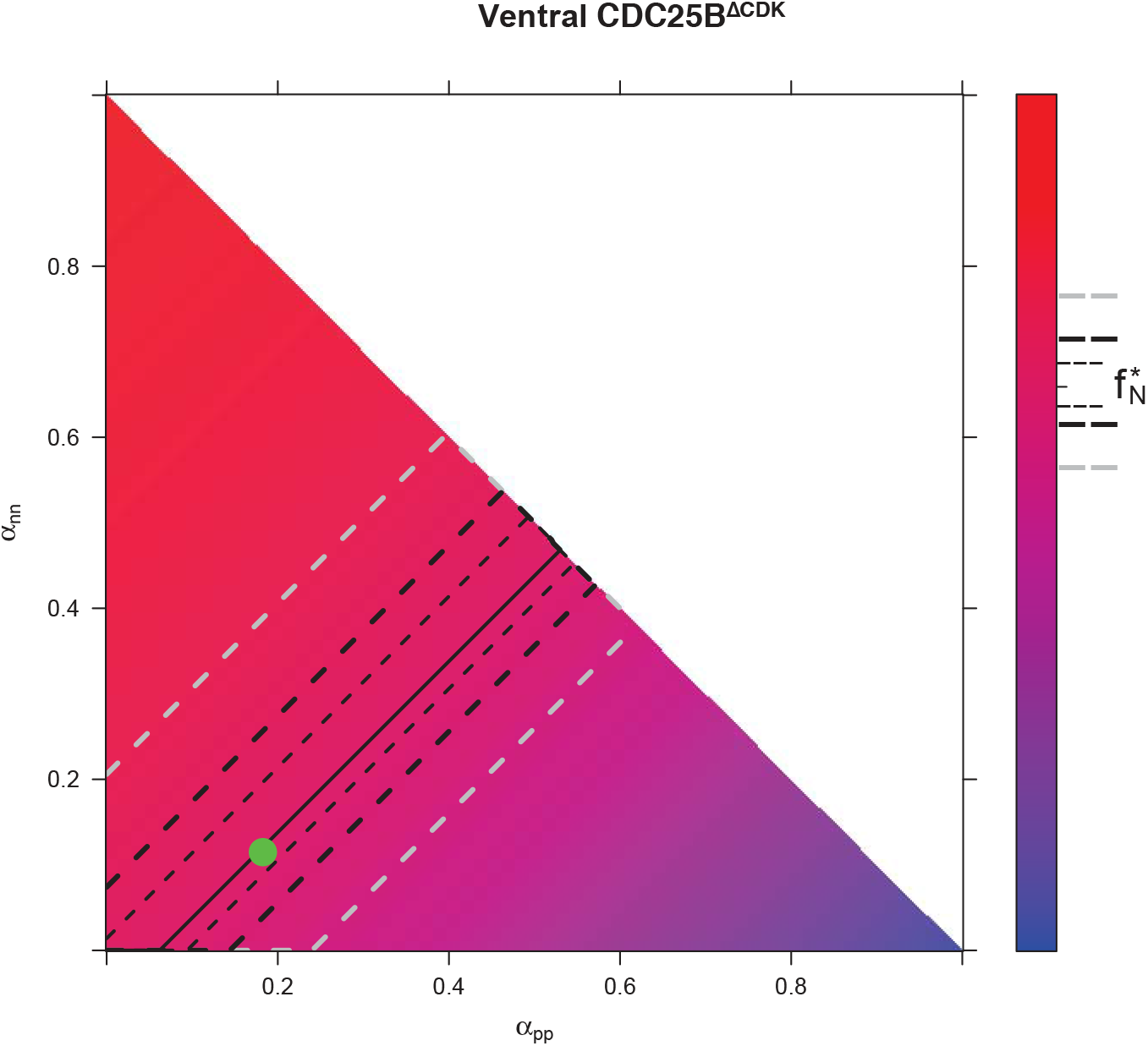
**Predicted** f_*N*_(48) **from** f_*N*_**(24) for every distribution of fates for** CDC25B^ΔCDK^ **condition in Ventral area**.

**Figure 8:**
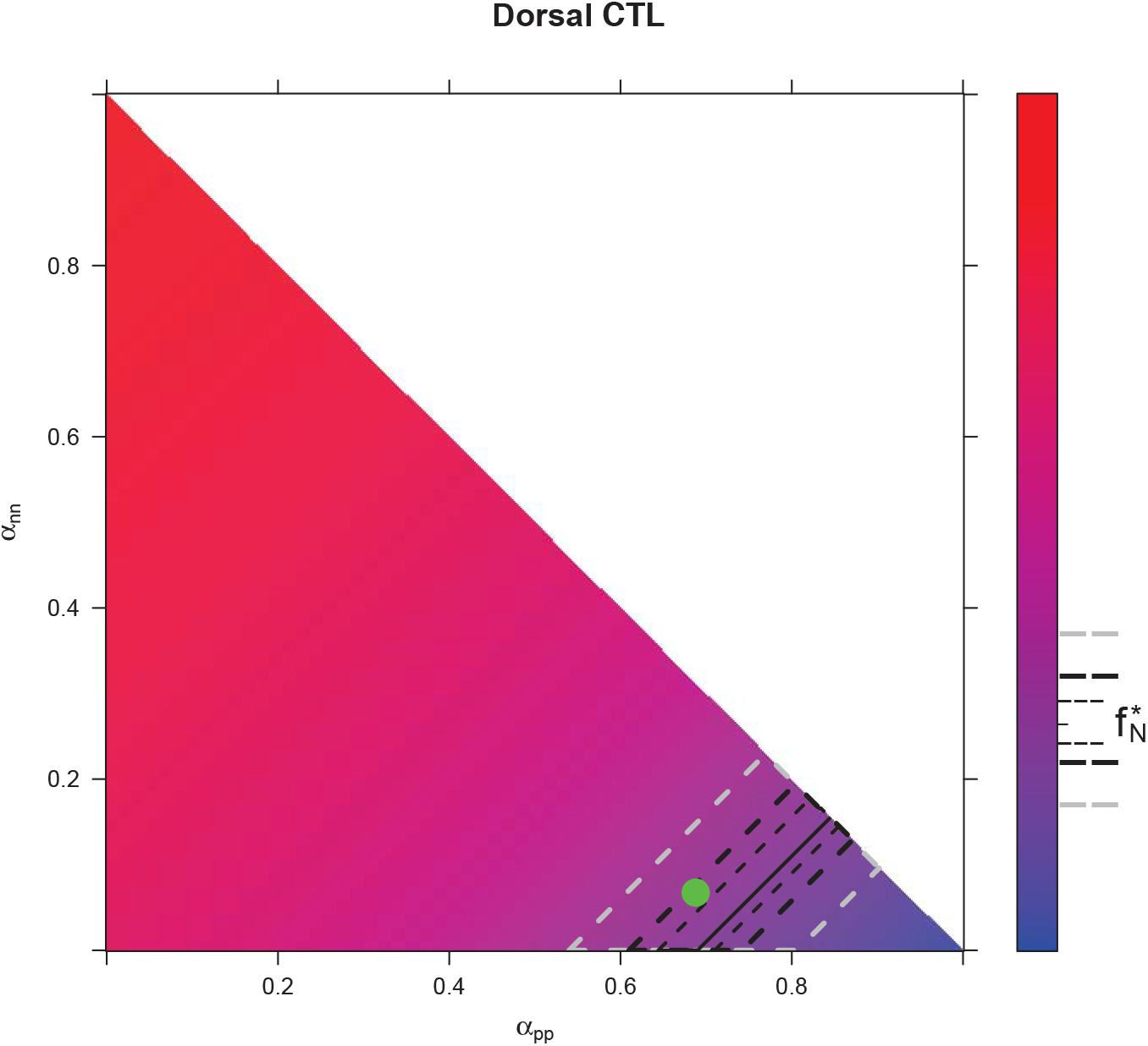
Predicted f_*N*_(48) from f_*N*_(24) for every distribution of fates for control condition in Dorsal area.

**Figure 9:**
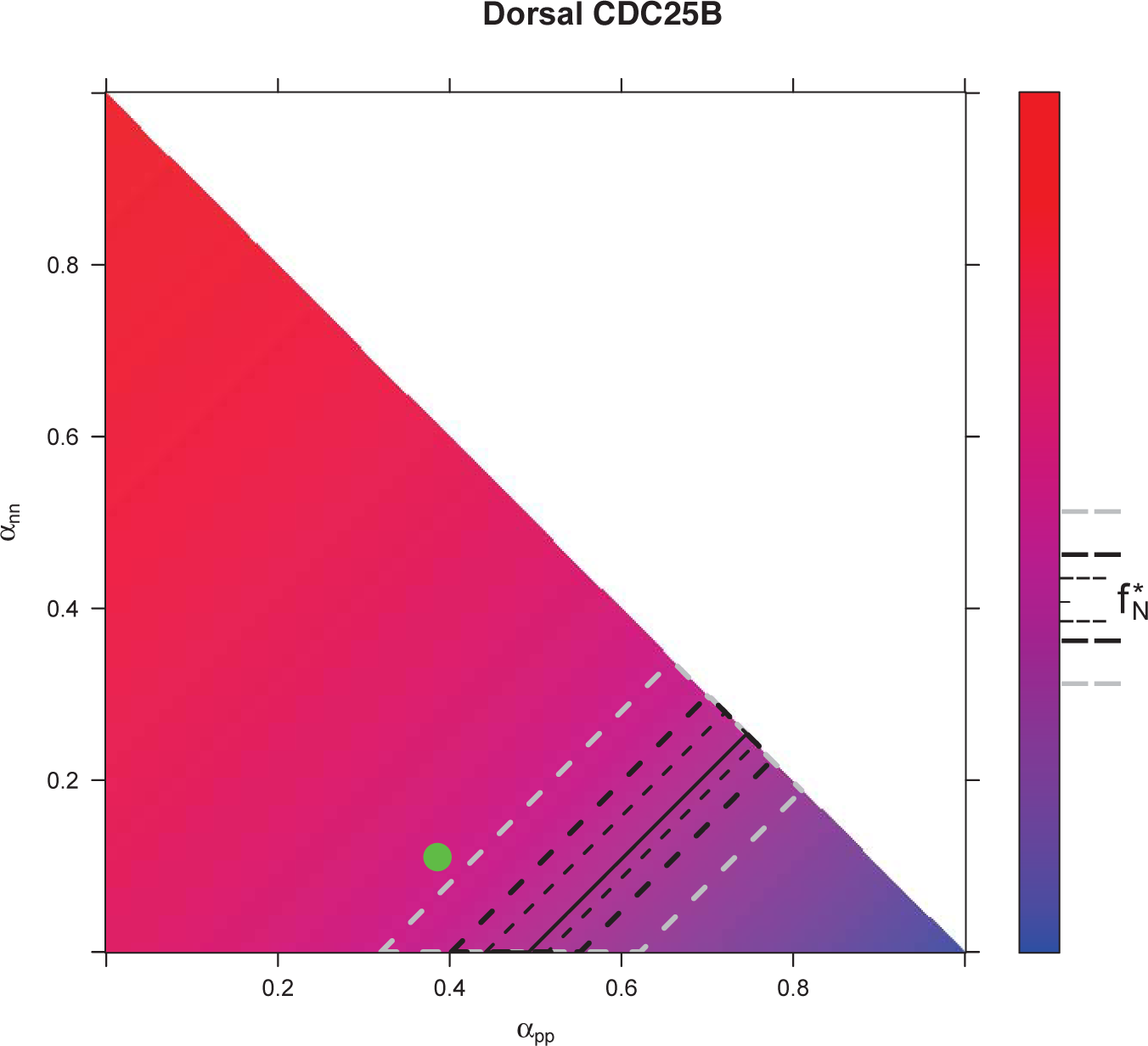
Predicted f_*N*_(48) from f_*N*_(24) for every distribution of fates for CDC25B condition in Dorsal area.

**Figure 10:**
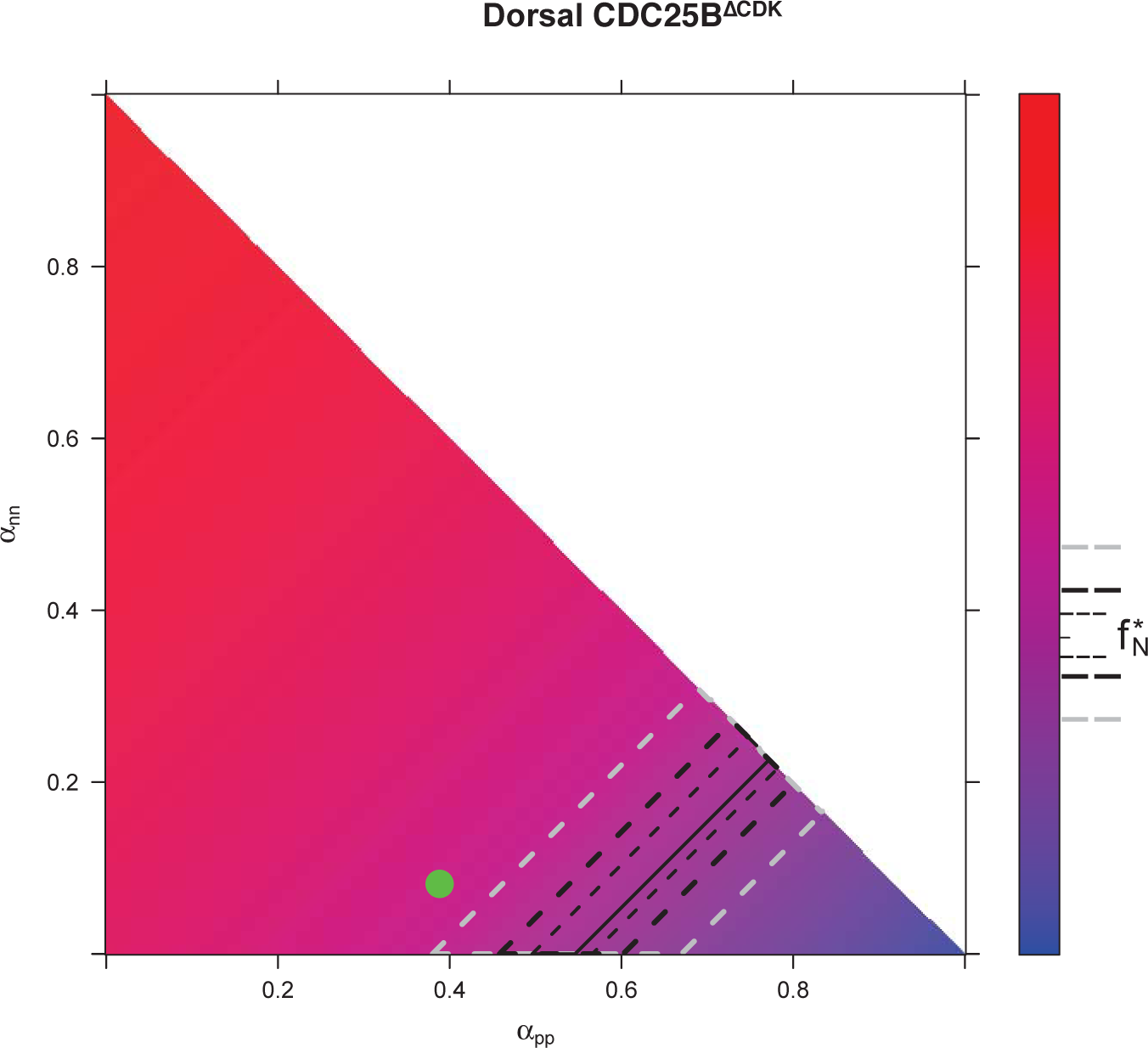
**Predicted** f_*N*_(48) **from f_*N*_(24) for every distribution of fates for** CDC25B^ΔCDK^ **condition in Dorsal area**.

#### 4.2 Predicted fraction of neurons at 48h knowing the fractions of neurons and the fate distribution at 24h

For computation of the predicted fractions of neurons at 48h (a.e.) reported in the main text (Figs. 6C), we used Eq.28, parametrized by the data obtained for the averaged fraction of neurons at 24h (a.e.), the fate distribution at 24h (a.e.), and the cell cycle 12h.

All predictions are gathered in fig. 11 as a function of the change in the balance proliferation/differentiation of the progenitors, induced by the CDC25B and the CDC25B^ΔCDK^ experiments. Together, the observations indicate that CDC25B and CDC25B^ΔCDK^ result in an increased proportion of neurons 48h a.e. (HH22).

**Figure 11:**
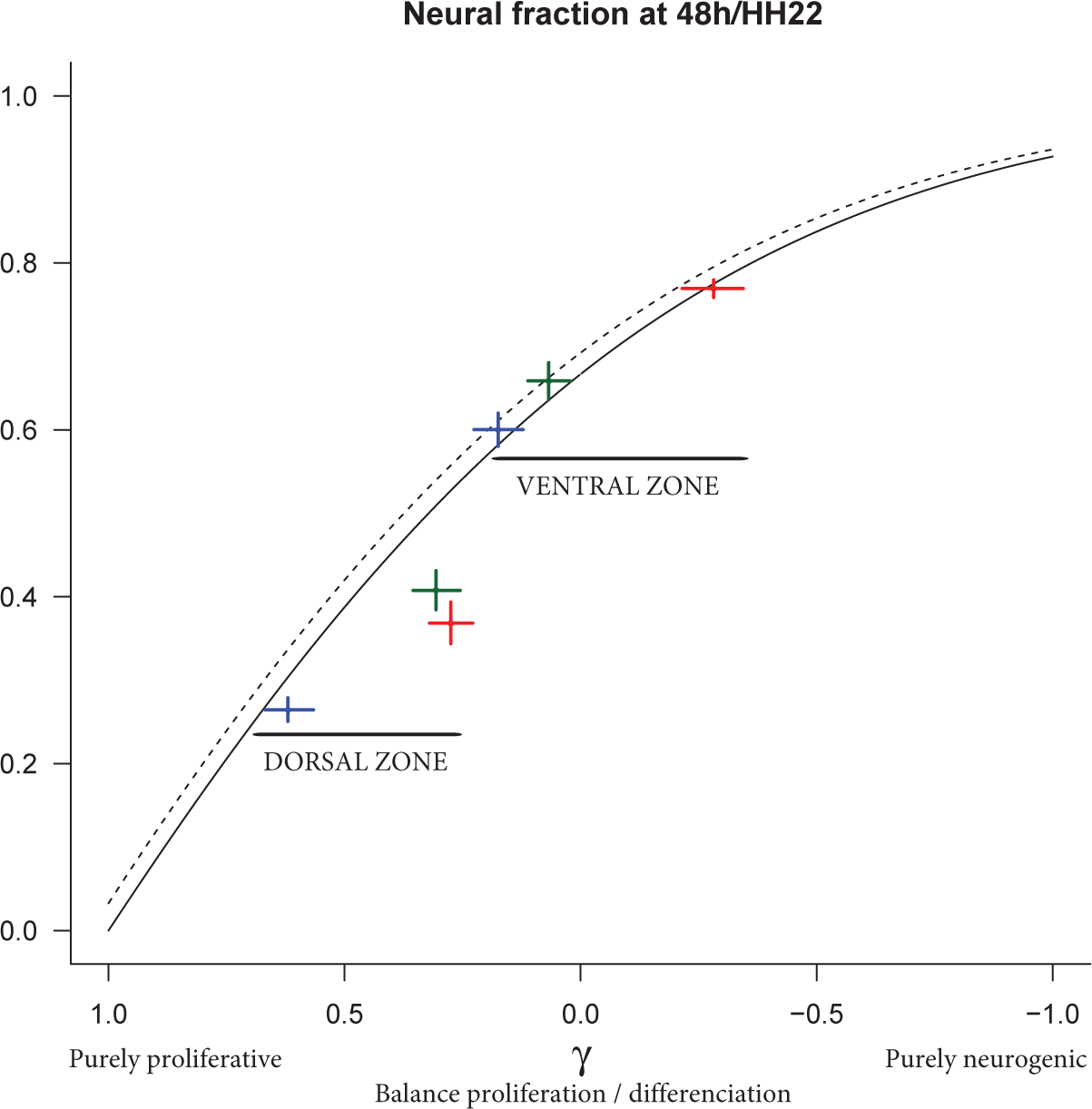
Predicted f_*N*_(48) from f_*N*_(24) varying the balance proliferation/differentiation *γ*. Plain line reports the model prediction for the dorsal zone, dotted line the model prediction for the ventral zone (predictions differ due to differences in the initial fraction f_*N*_(24) in the two zones). The experimental data are reported by crosses (cross arm length are 95% CI). Blue cross: CTL, red cross: CDC25B, green cross: CDC25B^ΔCDK^.

Such an increased *proportion* of neurons is actually compatible with two dynamical scenarios regarding how the absolute amounts of the two pools (progenitors, neurons) are modified by CDC25B gain of function: scenario 1) a speed-up of the neurons pool so that it increases faster under the gain of function at the expense of the progenitors pool expansion, or scenario 2) a decrease of the progenitors pool while the pool of neurons keeps the same expansion rate. Which scenario is relevant depends on how CDC25B affects the balance *γ* between proliferation and differentiation.

The pool of progenitors can increase only if *γ* > 0, which implies *α*_*pp*_ > *α*_*nn*_. In this case, the two pools can increase (scenario 1), their respective growth rates are controlled by *γ* and the neurogenic effect of CDC25B gain of function will produce a greater absolute number of neurons in the end (at 48h / HH22). Otherwise (*γ* < 0, i.e. *α*_*pp*_ < *α*_*nn*_), the pool of neurons can increase at about the same rate, yielding the same absolute number of neurons at 48h/HH22, and the increased fraction of neurons reflects a depletion of the pool of progenitors (scenario 2).

The model enlightens which is the most probable scenario for the dynamical impact of CDC25B manipulation since we can compute the underlying evolution of the absolute amounts of the two pools that determines the evolution of the neuronal fraction (Fig. 12C).

**Figure 12:**
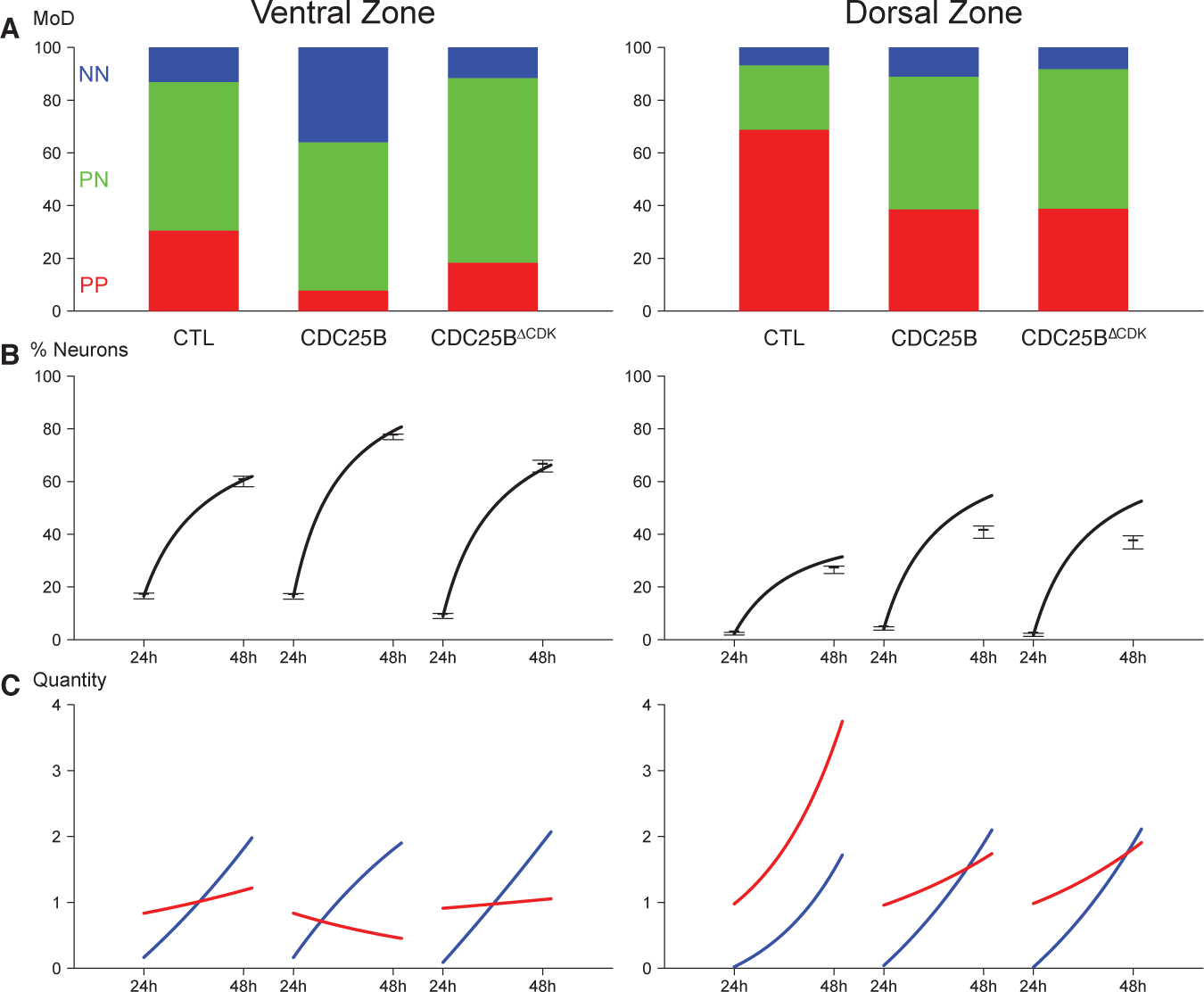
Summary of the data and predictions. A — Observed distributions of modes of divisions (MoD) for the three conditions and the two zone. B — Predicted evolutions of the neuronal fraction from f_*N*_(24) to f_*N*_(48) given the observed distribution of fates (lines) and observed fractions at 24h and 48h. C — Corresponding evolution in numbers of the two pools (Red: progenitors, Blue: neurons).

Under CDC25B gain of function in the dorsal neural tube (Fig. 12C-right), the percentage of progenitors performing pp-divisions keeps greater than the percentage of those performing nn-divisions (38.6% > 11.3%, *α*_*pp*_ > *α*_*nn*_) and the balance is still positive (*γ* = 0.386 − 0.113 = 0.273 > 0), so the pool of progenitors still increases but at a lower rate than control (where *γ* = 0.663 −0.078 = 0.585). The higher percentage of neurons at 48h/HH22 then results from an even higher absolute number of neurons (scenario 1).

By contrast, in the ventral neural tube, the balance shifts from *γ* = 0.393 −0.127 = 0.266 in control to *γ* = 0.069 - 0.407 =−0.338, becoming negative under CDC25B gain of function (scenario 2). Accordingly, the absolute number of neurons at 48h/HH22 is poorly affected, but the pool of progenitors declines, explaining the higher fraction of neurons (Fig. 12C-left).

